# A spatial atlas of human gastro-intestinal acute GVHD reveals epithelial and immune dynamics underlying disease pathophysiology

**DOI:** 10.1101/2024.09.02.610085

**Authors:** Nofar Azulay, Idan Milo, Yuval Bussi, Raz Ben Uri, Tal Keidar Haran, Michal Eldar, Ofer Elhanani, Yotam Harnik, Oran Yakubovsky, Ido Nachmany, Tomer-Meir Salame, Martin Wartenberg, Philippe Bertheau, David Michonneau, Gerard Socie, Leeat Keren

## Abstract

Acute graft-versus-host disease (aGVHD) is a significant complication of allogeneic hematopoietic stem cell transplantation (aHSCT), driven by alloreactive donor T cells in the gut. However, the roles of additional donor and host cells in this process are not fully understood. We conducted multiplexed imaging on 59 biopsies from patients with gastrointestinal GVHD and 10 healthy controls, revealing key pathological changes, including fibrosis, crypt alterations, loss of Paneth cells, accumulation of endocrine cells, and disrupted immune organization, particularly a reduction in IgA-secreting plasma cells. Interestingly, CD8T cells were enriched only in a subset of patients, while others exhibited non-canonical enrichments of macrophages and neutrophils. Post-transplantation time significantly influenced immune composition, with host cells dominating plasma and T cell compartments long after transplantation. This spatial atlas of healthy duodenum and GVHD uncovers non-canonical immune dynamics, offering insights into disease pathophysiology and potential clinical applications in GVHD and other inflammatory bowel diseases.

## Introduction

Worldwide, each year over 100,000 patients undergo an allogeneic hematopoietic stem cell transplantation (aHSCT) with the intent to cure high-risk hematologic malignancies, and non-malignant disorders ^1^. In aHSCT, the native immune system of the patient is submitted to a conditioning regimen with or without irradiation, and then reconstituted using hematopoietic stem cells that are transferred from a healthy donor. Healthy immune cells replace the malfunctioning immune and hematological systems and eradicate malignant tumor cells in a process called graft-versus-tumor (GVT). While aHSCT is an effective therapy for some patients, between 30%-70% of the patients will develop acute graft-versus-host disease (GVHD), a life-threatening complication of HSCT, in which immune cells from the donor attack organs in the recipient ^2^.

Among the different tissues affected by GVHD, gastrointestinal (GI) disease is the most difficult to treat and is the greatest cause of GVHD-related mortality ^2,3^. The small intestine has a distinct anatomical structure that facilitates the complex processes of digestion and absorption. The inner wall of the small intestine contains finger-shaped projections called villi intertwined with crevices that lead into the crypts of Lieberkühn. Different types of cells with diverse roles are present along the villi and crypts. Absorbing enterocytes, mucus-producing goblet cells and hormone-releasing endocrine cells form the villi, whereas stem cells and antimicrobial-releasing Paneth cells, form the crypts ^4,5^. Pathologically, GVHD is characterized by apoptosis of epithelial cells in low-grade disease, with increased cystically dilated crypts, villus atrophy and crypt destruction and loss in high-grade disease ^6^. Interestingly, pathological grades are only moderately correlated with the clinical grade, which is based on symptomatic disease severity ^7,8^. Recent work in mouse models suggested that the intestinal stem cells are the direct targets of attack in GVHD ^9^. However, information in humans is lacking, and while there has been some evidence regarding the fate of distinct epithelial subsets ^10–13^, a detailed description of epithelial cell fate in the gut across patients in GVHD is missing.

Much work has been dedicated to elucidating the pathophysiology of acute GVHD, demonstrating a major role for T cells in this process ^14,15^. Donor T cells recognize both major and minor histocompatibility antigens presented on host cells, leading to the activation and expansion of the T cells, and subsequent destruction of host cells. Indeed, the incidence of GVHD after HSCT increases with the number of mismatched human leukocyte antigen (HLA) alleles ^16^. Both CD4T and CD8T cells show tissue extravasation, clonal expansion and activated effector and tissue-resident phenotypes, which have been linked to the severity of the disease in murine models, non-human primates and human patients ^17–19^. The depletion of donor T cells results in a low incidence of acute and chronic GVHD, albeit at the expense of decreased GVT, leading to high rates of leukemia relapse^20^.

While alloreactive donor T cells are a driving factor in GVHD development, this simple model does not address many clinical observations. For example, GVHD is predominantly restricted to the GI tract, skin, and liver. Moreover, GVHD shows differential dynamics across individuals, with disease onset ranging from several days to several months post transplantation ^21^. Studies to investigate these observations have suggested the involvement of additional players in the pathophysiology of the disease, including distinct subsets of immune cells, epithelial cells of the gut and the microbiome ^22,23^. It was demonstrated that the conditioning therapy administered prior to HSCT can drive tissue damage, translocation of intestinal bacteria, changes in the composition of the microbiome, release of damage-associated and pathogen-associated molecular patterns and release of cytokines by resident cells in the tissue, which proceed the infiltration of donor T cells ^23–25^. Presence of monocyte-derived macrophages and neutrophils in the blood and affected tissues in human patients correlates with clinical severity and outcome ^26–28^, although their function was inconclusive in murine models ^29,30^. Moreover, recent evidence suggests that tissue-resident immune cells of the host, both T cells and myeloid cells, may survive the conditioning regimens and play an important role in instigating tissue inflammation ^31–34^. However, the interactions of these distinct donor and host immune populations and their role in disease progression remains unclear. It is also not known if these distinct cell types are equally important across patients, and what is their relative contribution to distinct stages of the disease.

Here, we leveraged Multiplexed Ion Beam Imaging by Time of Flight (MIBI-TOF) ^35,36^, to generate comprehensive spatial maps of healthy and GVHD-afflicted duodenum at single cell resolution. We profiled 59 biopsies taken from GVHD patients and 10 healthy controls, and measured the spatial expression of 40 proteins, designated to delineate distinct epithelial cell populations of the gut, stromal cells and immune cells. Normal duodenum was highly stereotypical across individuals, showing high organization and multiple layers of zonation within epithelial, stromal and immune cells. In contrast, GVHD manifests in increased fibrosis, alterations in crypt morphology, loss of protein zonation and altered composition of distinct epithelial subsets. We observed a reduction in Paneth cells, and an increase in abnormal crypt morphologies enriched with endocrine cells. The immune composition in GVHD was characterized by a breakdown of homeostatic composition and organization, with a major reduction in IgA-secreting plasma cells. Interestingly, CD8T cells dominated the immune landscape in only a subset of patients, whereas others could be classified into subtypes characterized by non-canonical enrichments of other immune cell types. In particular, the macrophage-enriched subtype associated with poor clinical grade and low survival. Time post-transplantation also significantly influenced the composition of the cellular infiltrate, showing shared dynamics among individuals. Using fluorescent in situ hybridization (FISH) in sex-mismatched cases, we distinguished between donor and host immune cells. Host cells predominated in the plasma and T cell populations in the gut for prolonged periods after transplantation, suggesting that other factors beyond donor-derived T cells may contribute to GVHD variability. We also identified intra-patient variability in disease severity, with highly localized immune responses, at times eliminating a single crypt while leaving the neighbouring crypt intact. These results highlight the importance of fine-grade spatial analysis that can uncover short-scale immune interactions and suggests local inflammation as a defining driving force of disease progression. Overall, this spatial atlas of healthy duodenum and GVHD reveals non-canonical immune dynamics underlying disease pathophysiology in humans with potential implications for clinical diagnosis and therapeutics in both GVHD as well as other inflammatory bowel diseases.

## Results

### A spatial atlas of healthy and aGVHD duodenum in human patients

To construct a spatial atlas of aGVHD we assembled a retrospective cohort of duodenal (N=40) and colonic (N=19) biopsies taken from patients who underwent aHSCT and were clinically suspected to develop aGVHD (**Fig. 1A, Table. S1**). Thirty-four patients were diagnosed with mild clinical GVHD in the GI tract (stages 0 and 1) while 25 manifested severe clinical symptoms (stages 2 and 3). After pathological examination, 43 were diagnosed with mild pathological GVHD (stages 0 and 1) and 16 with severe pathological GVHD (stages 2 and 3). Following diagnosis, patients were treated with corticosteroids, to which 29 responded, and 23 were refractory (**Table. S1**). In this cohort, anatomical location was associated with severity of aGVHD, with colonic biopsies belonging to patients with higher pathological scores and cortico-refractory disease (**Fig. 1A).** In addition, we collected specimens of ten healthy duodenums taken from Whipple procedures to serve as control.

**Figure 1.**
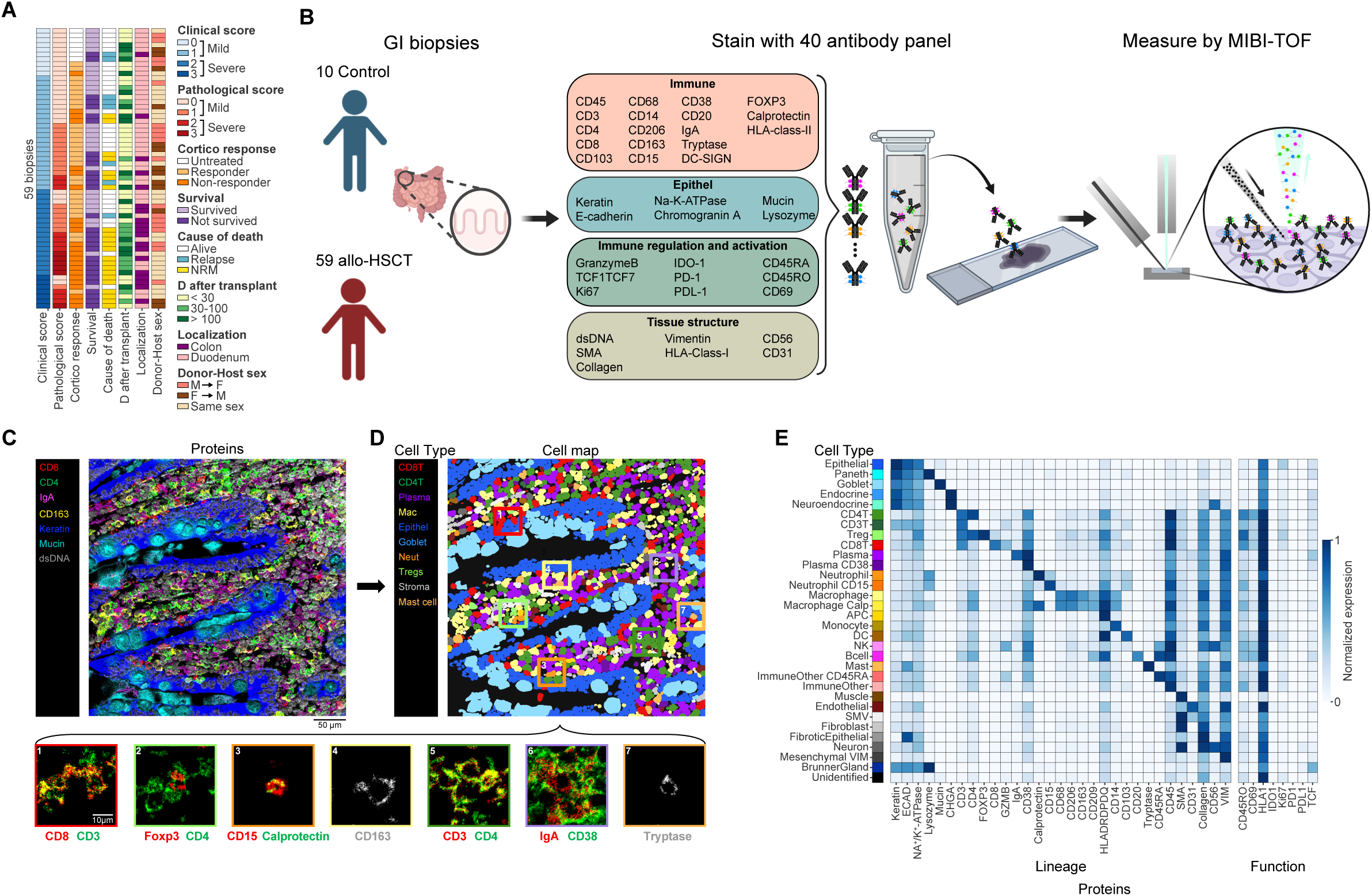
Multiplexed imaging of healthy and diseased duodenum reveals distinct cell populations. **(A)** Clinical characteristics of the GVHD cohort. Rows are sorted in a hierarchical manner from the left to right columns. NRM - Non relapse mortality. **(B)** Clinical tissue specimens were stained using a mixture of antibodies labeled with elemental mass tags and measured by MIBI-TOF to produce 40-dimensional images depicting protein expression and localization. **(C)** MIBI-TOF staining of seven proteins in one field of view (FOV). **(D)** Segmented and classified cells in the same FOV shown in C. Bottom; zoom-in on specific cells showing the expression of canonical markers. **(E)** Heatmap depicting the mean expression level of each protein (x-axis) in each cell type (y-axis). Left: lineage proteins used for classification. Right: Phenotypic proteins not used for classification.

To study how immune composition and organization relates to disease progression and outcome, we designed a panel of forty antibodies to interrogate the gut-immune microenvironment (**Fig. 1B, Table. S2**). The panel included antibodies to identify populations of the gut epithelium, stromal cells, and immune cells. We validated the specificity and sensitivity of all antibodies on control tissues (**Fig. S1)**. We used this panel to stain formalin fixed paraffin embedded (FFPE) tissue specimens from our cohort and imaged them using MIBI-TOF ^36^. Overall, we acquired 263 high-dimensional images, each depicting the spatial expression of forty proteins *in situ* (**Fig. 1C**). To map single cell identity and function we segmented individual cells using our previously validated deep-learning pipeline ^37^, with modifications to account for large goblet cells (methods). We then classified individual cells using CellTune, a human-in-the-loop machine learning framework for cell classification (methods). Altogether, we identified 32 cell populations, including four populations of epithelial cells (enterocytes, goblet, endocrine, and Paneth cells), various immune populations (including CD4T and CD8T cells, Tregs, plasma, B cells, neutrophils, macrophages and mast cells) and several stromal populations (including fibroblasts, endothelial cells, muscles, neurons), well-aligned with the literature of expected cell types in the duodenum (**Fig. 1D, E)** ^38^.

### The immune landscape of healthy duodenum is characterized by a stereotypic zonated organization across individuals

To generate a baseline of healthy duodenum, we first analyzed the images from the controls, and added four samples that were specifically embedded and cut to expose the entire axis of the villi from its bottom to the tip. We identified the unique microanatomical structure of the small intestine: the submucosal layer stained by collagen, the muscularis mucosa stained by SMA, and the mucosal layer comprised of the epithelium and the lamina propria (**Fig. 2A**). As expected, we observed specific localizations of distinct epithelial cells along the crypt-villus axis, shared across individuals ^39,40^. Proliferating Ki-67+ epithelial cells were located in the base of the crypt, near anti-microbial peptide secreting Paneth cells (**Fig. 2B**). These proliferating cells at the base will differentiate into mature epithelial cells that migrate up the crypt-villus axis ^5^. Mucus-secreting goblet cells and hormone-secreting endocrine cells were equally spaced along the villus. Across individuals, we found a relatively homogenous distribution of cell populations (STD = 0.01, **Fig. 2C**). In accordance with previous reports ^40^, epithelial populations generally constitute almost 40% of the total number of cells in the image, ranging from high prevalence (enterocytes 32%) to low prevalence (goblet 4%, Paneth and endocrine <1%). Immune cells were highly abundant and mainly found in the lamina propria (40%).

**Figure 2:**
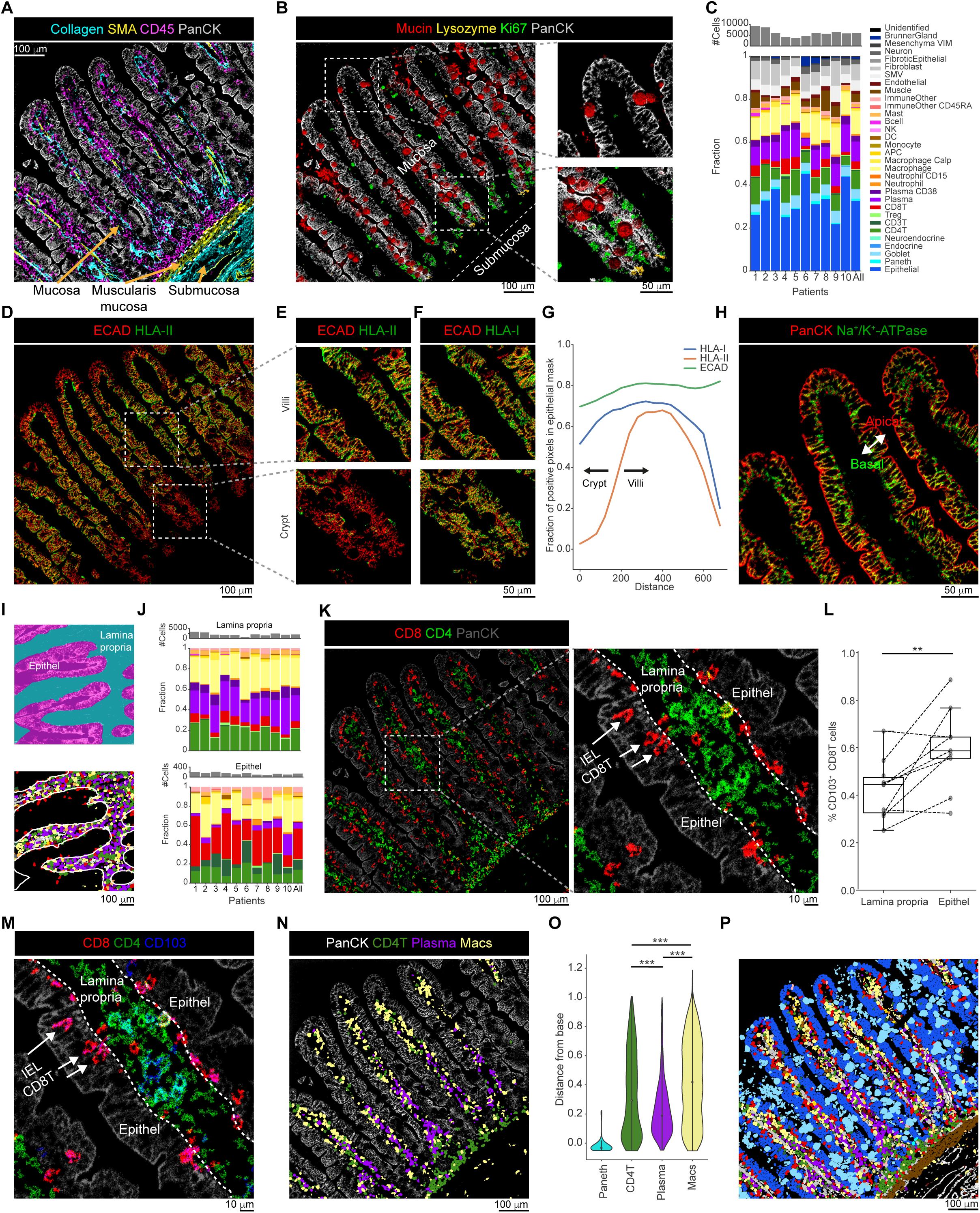
Stereotypical organization in healthy duodenum. **(A)** MIBI-TOF image of healthy duodenum shows the microanatomical structure of the small intestine. **(B)** Location of epithelial cells in the duodenum. Paneth cells are located at the base of the crypts, next to amplifying cells. Goblet cells are scattered along the Villus. **(C)** Top: Number of cells (y-axis) across 10 healthy individuals (x-axis). Bottom: Cell composition (y-axis) across 10 healthy individuals (x-axis). **(D)** MIBI-TOF image showing the expression of E-cadherin (ECAD, red) and HLA-DR-DP-DQ (HLA-II, green) in epithelial cells. Expression is shown only in the epitelial mask. **(E, F)** Magnified images of the two insets shown in D depicting expression of HLA-II (E) and HLA-I (F). **(G)** Shown is the expression (y-axis) of HLA-I, HLA-II and E-cadherin (ECAD) along the crypt-villus axis (x-axis) for the image in (D). HLA-II expression is absent in the base of the crypts. **(H)** MIBI-TOF image showing basoapical expression of Pan-keratin (PanCK, red) and Na+/K+-ATPase (green). **(I)** Top: Pixel masks to stratify regions inside (pink) and outside (cyan) the epithelium. Bottom: Shown are the immune cells in the same FOV. The white line indicates the borders of the epithelium. Cell types are colored as in (C) **(J)** Immune composition (y-axis) across patients (x-axis) inside (bottom) and outside (top) the epithelium. Bar colors are as in (C). **(K)** MIBI-TOF Image showing the expression of CD4 (green), CD8 (red) and Pan keratin (PanCK, gray). Right: Magnified inset. Dashed lines demarcate the borders of the lamina propria. CD4T cells mostly reside in the lamina propria while CD8T cells reside both in the lamina propria and inside the epithelium. Intraepithelial lymphocytes (IEL). **(L)** Percentage of CD103+ cells out of the CD8T cells (y-axis) inside and outside the epithelium. Each dot represents one patient (n = 9, patients with less than 10 cells in the epithelium were excluded from the analysis). ** P<0.01, paired t-test. **(M)** MIBI-TOF Image showing the expression of CD4 (green), CD8 (red) and CD103 (blue). Dashed lines demarcate the borders of the lamina propria. IEL CD8T cells are preferentially CD103+. **(N)** Representative FOV demonstrates zonation of CD4T cells (green), plasma cells (purple) and macrophages (yellow). **(O)** Shown are the distances from the base of the crypt (y-axis) for different cell types (x-axis) in four controls. *** P<0.0001, Wilcoxon rank-sum test followed by Benjamini-Hochberg adjustment. **(P)** The same image as in (N) with all cell types. Cell types are colored as in (C).

Recent studies showed that the intestinal epithelium is functionally organized, with gene expression gradients both along and perpendicular to the villi-crypt axis ^41,42^. We therefore evaluated the expression of different proteins along the major and minor axes (methods). We found two proteins with zonated expression patterns in epithelial cells. The first was HLA class II (HLA-II), which consistently showed expression in the villi, which reduced towards the base and was completely absent in the crypts. We also observed a reduction of both HLA class I (HLA-I) and HLA-II expression at the tip of the villi in some but not all patients. (**Fig. 2D-G, S2A**). While the expression of HLA-II on GI epithelial cells has been suggested to play a role in antigen expression and regulation of gut immunological homeostasis; the functional role of its zonation in both mouse and human remains unclear ^42–45^. In addition, we found a baso-apical polarization of Na^+^/K^+^-ATPase, which was predominantly expressed in the basolateral side of the epithelium (**Fig. 2H**). Na^+^/K^+^-ATPase maintains the transcellular Na+ gradient which is essential to facilitate Na-solute co-transport processes of intestinal epithelial cells ^46^, and as such its basolateral positioning next to blood vessels may reflect intra-cellular spatial specialization.

Next, we interrogated the composition and spatial distribution of immune cells in normal duodenum. We found that plasma, macrophage and CD4T were the most frequent cell types, with prevalences of 30%, 27% and 20%, respectively. T regulatory cells (Tregs), neutrophils and other immune cells had low prevalence (< 2% each), and CD8T (8%) fell in between. As expected, we hardly observed B cells among the normal samples, presumably because we focused on the mucosal layer, where B cells are scarce ^4^ (**Fig. 2C**). We explored how different immune cells organize relative to epithelial cells by using a pixel classifier to stratify our images into two regions, inside and outside the epithelium (methods, **Fig. 2I**). We confirmed that immune composition differed between the lamina propria and intra-epithelial cells. The lamina propria was enriched with plasma cells, macrophages and CD4T cells, whereas CD8T and DN T cells (CD3T cells, double negative for CD4 and CD8) were the predominant populations inside the epithelium (**Fig. 2J, K, S2B**). These intraepithelial lymphocytes (IELs) expressed high levels of CD103, indicating their residency phenotype (paired t-test p=0.009 **Fig. 2L, M**). Resident IELs may play an important role in gut immunity by killing infected epithelial cells and thus protecting the intestine from pathogens ^47^.

Finally, we investigated immune zonation along the crypt-villi axis. We found that CD4T cells mainly accumulated in the base but also resided in the upper part of the villi. In contrast, plasma cells populated the region right above the crypts, while being scarce at the base or the tip of the villi. The upper part of the villi was predominantly occupied by antigen-presenting macrophages (**Fig. 2N-P**). This organization was highly reproducible across individuals (**Fig. S2C**), suggesting a universal zonation of immune populations in the gut.

### GVHD is characterized by a disrupted epithelium and increased fibrosis

We explored how GVHD affects the structure of the gut epithelium. To this end, we adapted Cellpose, a deep-learning cell segmentation model ^48^, to identify individual crypts in our images (**Fig. 3A, Fig. S2D**, methods). In the control group, the crypts had a relatively narrow size distribution, whereas in GVHD samples we identified both very small and very large crypts. Larger crypts were mainly from the colon, whereas smaller crypts were mostly associated with higher pathological grade in the duodenum (**Fig. S2E**, **Fig. 3B**). We quantified the distance between neighboring crypts (methods) and found that in the controls the crypts were regularly spaced, whereas in GVHD the crypts showed a larger variance in their spacing, with a significant subset of isolated crypts (**Fig. S2F**). This isolation correlated with the pathological stage of the disease, with higher stage disease showing more isolated crypts (**Fig. 3C**). Accordingly, a quantification of cell types across GVHD patients showed high degree of variability in epithelial cells with many patients exhibiting a major decrease, which was also accompanied by an increase in fibroblasts and correlated with high-grade disease (**Fig. 3D, E**). Altogether, in GVHD the global structure of the crypt morphology is altered and characterized by increased fibrosis (**Fig. 3F**).

**Figure 3:**
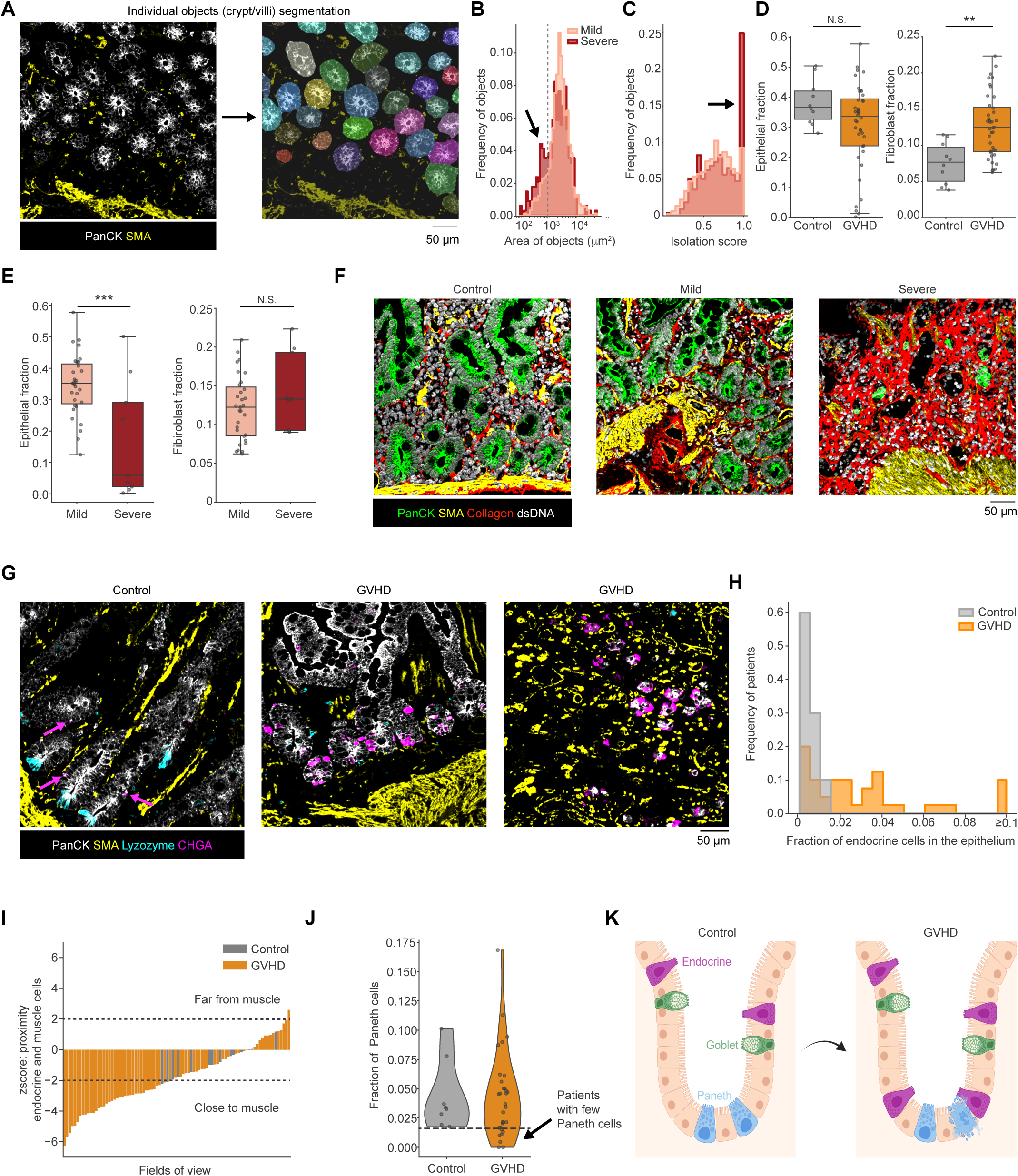
GVHD is characterized by disrupted epithelium and increased fibrosis. **(A)** Images of epithelial proteins (left) were used to segment individual crypt/villus objects (right) using Cellpose. **(B)** Histograms of the distribution of crypt/villi sizes in patients with GVHD diagnosed as pathologically mild (pink) and severe (red). Arrow denotes accumulation of small crypts in severe GVHD. **(C)** For each crypt/villus an isolation score was calculated as the percentage of the object’s non-overlapping area with neighboring crypts within a 75-pixel expansion. Histograms show the distribution of the isolation scores in patients with GVHD diagnosed as pathologically mild (pink) and severe (red). Arrow denotes accumulation of isolated crypts in severe GVHD. **(D)** Fraction of epithelial cells (left) and fibroblasts (right) out of all cells in control (gray) and GVHD patients (orange). **(E)** Fraction of epithelial cells (left) and fibroblasts (right) out of all cells in pathologically mild (pink) and severe patients (red). Statistical tests were conducted for (D,E) by Wilcoxon rank-sum test, **P<0.01, ***P<0.001 (Control n = 10, Duodenum GVHD n = 40). **(F)** Images demonstrating the loss of epithelial cells in GVHD and the development of fibrosis. **(G)** Representative images displaying the sparsity of endocrine cells (CHGA, magenta, indicated by arrows) in control (left image) versus the hyperplasia in GVHD, along with the loss of Paneth cells (lysosyme, cyan). **(H)** Histogram of the fraction of endocrine and neuoroendcrine cells out of epithelial cells in control and GVHD patients. **(I)** The mean distance of the centroids of endocrine and neuoroendcrine cells to the muscle mask was calculated and tested against a baseline distribution generated by permuting the location of these cells in the epithelium. Shown is the z-score of enrichment of endocrine cells next to the muscle (y-axis) across FOVs (x-axis). FOVs are ordered according to z-score. Dashed lines indicate a difference of 2 standard deviations from the mean. Negative values indicate that the endocrine cells are enriched closer to the muscle. Patients without muscle regions were excluded from the analysis. Control n = 8 and duodeneal GVHD n = 27. **(J)** Violin plot of the frequency of Paneth cells in control and GVHD patients. To examine physiologically-relevant regions, patients without muscle regions were excluded from the analysis resulting with control n = 9 and duodeneal GVHD n = 31. **(K)** Schematic summarizing the loss of Paneth cells and accumulation of endocrine cells in GVHD patients.

Next, we gauged the fate of different epithelial cells in GVHD. First, we evaluated endocrine cells. We found that many GVHD patients displayed an increase in the proportion of endocrine cells, perhaps indicating their selective sparing in GVHD ^11,12^ (**Fig. 3G, H**). Moreover, we identified a recurring structure of small, isolated crypts, with a high density of endocrine cells and high expression of Chromogranin A (**Fig. 3G, Fig. S2G**). To evaluate the location of the endocrine cells, we trained a pixel classifier to identify regions of the muscularis mucosa and calculated the distances between the endocrine cells and the muscularis mucosa. We then compared these distances to the baseline distances of the epithelium by permuting their locations within the epithelium (methods). While in normal duodenum the endocrine cells were evenly spread along the villi, GVHD showed increased density primarily near the muscularis mucosa (**Fig. 3G, I**). Next, we evaluated Paneth cells. In accordance with previous reports ^49^, we found a reduction in Paneth cells in a subset of patients, but not in all (**Fig. 3J**). Since Paneth cells are localized in the base of the crypts, we verified that this reduction is not an artifact of sampling distinct regions in different patients. To this end, we only analyzed fields of view in which the muscularis regions were sampled (methods). Our results support clinical observations showing an increase in the anti-microbial protein REG3A in the plasma of GVHD patients, presumably due to the destruction of Paneth cells ^50,51^. Altogether, these results may suggest that GVHD drives a process whereby Paneth cells at the base of the crypts are lost and replaced by abnormally differentiated endocrine cells (**Fig. 3K**).

### Spatial mapping of human aGVHD reveals non-canonical immunological subtypes

To investigate the local immunological processes in GVHD we analyzed the immune composition across GVHD patients and controls. We found that GVHD was characterized by a significant decrease in plasma and CD4T cells and an increase in various stromal cells, mainly fibroblasts, in agreement with previous observations^19^ (**Fig. 4A**). Comparing between patients with mild and severe clinical manifestations further enhanced these trends and revealed infiltration of additional immune cells in the severe patients including neutrophils, Tregs, macrophages and CD8T cells (**Fig. 4A, B**). To evaluate the consistency of these trends across patients, we clustered all fields of view (FOVs) (**Fig. 4C**) and patients (**Fig. S3A**) according to their immune composition. We found that samples from healthy duodenum clustered together, presenting similar compositions (±30%) of plasma cells, CD4T cells and myeloid cells. Some GVHD samples displayed a normal-like immune composition and clustered with healthy duodenum. Others showed a breakdown of this organization, clustering into distinct immunological subtypes (**Fig. 4C, D, S3A-D**). The different subtypes were characterized by enrichments of distinct cell types, including CD8T cells, neutrophils and macrophages. We found that some subtypes were associated with distinct clinical parameters. For example, the Macs + CD8T subgroup was associated with decreases in epithelium accompanied by increases in stroma and overall worse clinical scores (**Fig. 4C**, red highlight and **Fig. S3E,** p=0.003). The CD8T-enriched subgroup associated with later times after transplantation (**Fig. 4C**, green highlight and **Fig. S3F**, p=0.001). Overall, we conclude that although CD8T cells play a prominent role in GVHD ^19^, the disease involves additional immune cells, with differential impact across patients.

**Figure 4:**
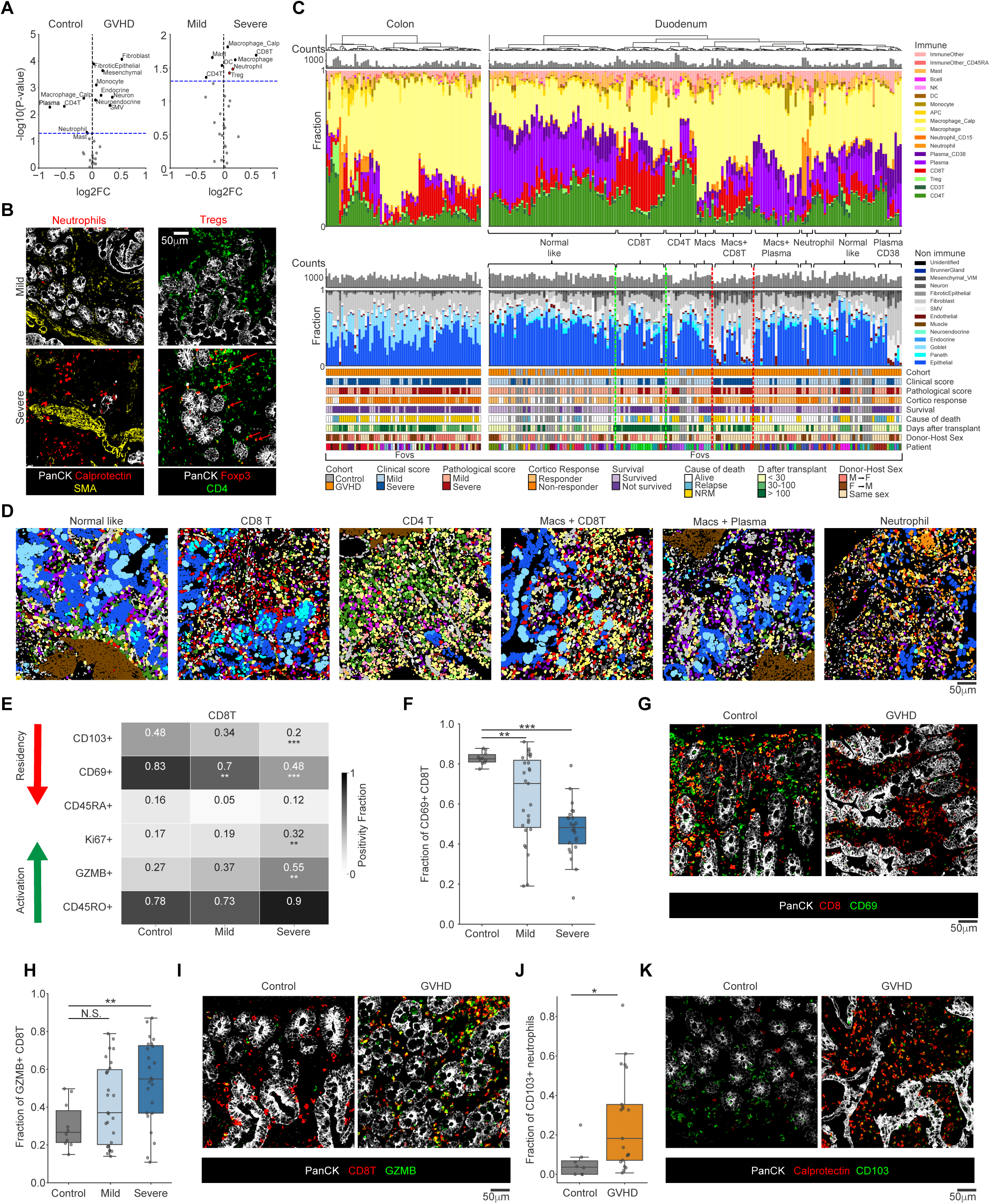
Spatial mapping of human aGVHD reveals non-canonical immunological subtypes. **(A)** Volcano plots showing differences in cell density in control and GVHD (Left) or clinically mild and severe patients (Right). Blue dashed line marks p-value=0.05. Black: significantly enriched cell types. Red: Example images are shown in B (control n = 10, duodenum GVHD n = 40). **(B)** Example images of neutrophils (left) and Tregs (right) in clinically mild (top) and severe (bottom) patients. **(C)** Top: Total number of immune cells and their relative abundances (y-axis) across all FOVs (x-axis) in samples from the colon (left) and duodenum (right). Bottom: The same for non-immune cells. Below are shown clinical characteristics of the patients. Red dashed lines highlight the group of Macs+CD8T associated with poor clinical score (Chi-square p=0.003). Green dashed lines highlight the group of CD8T associated with later time after transplantation (p=0.001). NRM: Non-relapse mortality. **(D)** Representative cell map images of different immune subtypes. **(E)** For each protein (rows), shown is the fraction of positive cells out of the CD8T cells in control, clinically mild and clinically severe GVHD (columns). Patients with more than 20 CD8T cells were included (control n = 10, GVHD duodenum n = 33, GVHD colon n = 18). Asterisks denote statistically significant changes relative to the control **(F)** Fraction of CD69+ cells out of the CD8T cells (y-axis) in control, clinically mild and clinically severe patients (x-axis). Each dot represents one patient. **(G)** Images showing the expression of CD69 (green) in CD8T cells (red) in GVHD (right) compared to control (left). **(H)** Fraction of GZMB+ cells out of the CD8T cells (y-axis) in control, clinically mild and clinically severe patients (x-axis). Each dot represents one patient. **(I)** Images showing the expression of GZMB (green) in CD8T cells (red) in GVHD compared to control. **(J)** Fraction of CD103+ cells out of neutrophils (y-axis) in GVHD compared to control (x-axis). Patients with more than 20 neutrophils were included (control n = 8, GVHD duodenum n = 12, GVHD colon n = 9). **(K)** Images showing the expression of CD103 (green) on neutrophils (calprotectin, red) in GVHD compared to control. Statistical tests for in this figure were conducted by Wilcoxon rank-sum test followed by Benjamini-Hochberg p-value adjustment, *P<0.05, **P<0.01, ***P<0.001, unless otherwise indicated.

Next, we explored phenotypic and cell state changes in immune cells associated with GVHD (**Fig. 4E, S3G-K**). We examined the expression of seven functional proteins in the T cell compartment. For each protein, we quantified the fraction of positive cells in each patient and tested for significant changes between healthy controls, mild and severe GVHD. We found that in GVHD both CD4T and CD8T cells lost their expression of CD103 and CD69, with more pronounced reductions observed for severe GVHD patients (**Fig. 4E-G, S3G)**. For example, the percent of CD69+ CD8T cells dropped from 83% in healthy controls to 48% in severe GVHD patients (p<0.001). Together, this suggests a loss of tissue-resident cells. This loss could be a direct effect of the conditioning regimen given prior to HSCT; or could be attributed to the entrance of new cells from the blood, likely originating from the donor. Concomitantly with the reduction in naïve and residency phenotypes, the CD8T cells acquired an activated and proliferating phenotype, as manifested by a ±2-fold increase in the fraction of GZMB+ CD8T cells (p<0.01, **Fig. 4E, H-I**).

In contrast to T cells, neutrophils displayed significantly elevated levels of CD103 in GVHD, with a 5-fold increase between control and GVHD patients (p=0.02, **Fig. 4J, K, S3K**). CD103 is an integrin that engages with epithelial cells and has been associated with the maintenance of tissue resident lymphocytes. Its expression on neutrophils may suggest that a residency phenotype emerges in neutrophils in GVHD. This hypothesis is further supported by the correlation between the expression of CD103 on neutrophils and their number in the tissue (Pearson R=0.58, p=0.005, **Fig. S3L**). However, since expression of CD103 on neutrophils has not been studied extensively yet, it is possible that CD103 plays a role in other cellular processes which are not related to tissue residency and maintenance. Altogether, we found that GVHD, and especially clinically severe GVHD, is characterized by a breakdown of immune homeostasis, with a universal reduction in plasma cells, and increase in distinct immune cells, including macrophages, neutrophils and T cells. These changes in composition further associate with functional changes in both lymphocytic and myeloid compartments.

### Temporal kinetics of immune cells of host and donor origin show persistent host T cells and plasma cells following transplantation

Our analysis of immune cell composition showed that the CD8T enriched subgroup associated with later times after transplantation (**Fig. 4C**). We therefore hypothesized that the heterogeneity in immune populations across patients may be partly driven by the time in which the biopsy was taken relative to the transplantation, a parameter that is frequently overlooked. To this end, we sorted all patients according to the time after transplantation and evaluated the frequency of each immune cell type. Interestingly, we found that the prevalence of distinct immune populations was correlated with the time after transplantation. Across patients the fraction of plasma cells and mast cells dropped between 60% to 90% within the first month after transplantation. This drop was accompanied by a compensatory, yet slower increase in macrophages, neutrophils and Tregs, which all peaked between 50 to 100 days post transplantation. Finally, CD8T cells, often regarded as the instigators of the disease increased in frequency much later, peaking months after transplantation (**Fig. 5A, B**). Although our cohort is not powered to examine very late timepoints after transplantation, anecdotal results from two patients whose biopsies were taken more than a year after transplantation suggest that this disruption of immune homeostasis may not return to normal levels even several years after transplantation (**Fig. 5A**, plasma cells). Alternatively, this could be a unique manifestation of late-onset acute GVHD ^52^, or the result of treatment with steroids before taking the biopsy (**Supp. Table 1**). Overall, this analysis suggests stereotypical immune kinetics in the gut following HSCT, and points to early events, such as the decrease in plasma cells and mast cells as important factors in the initiation of the disease.

**Figure 5:**
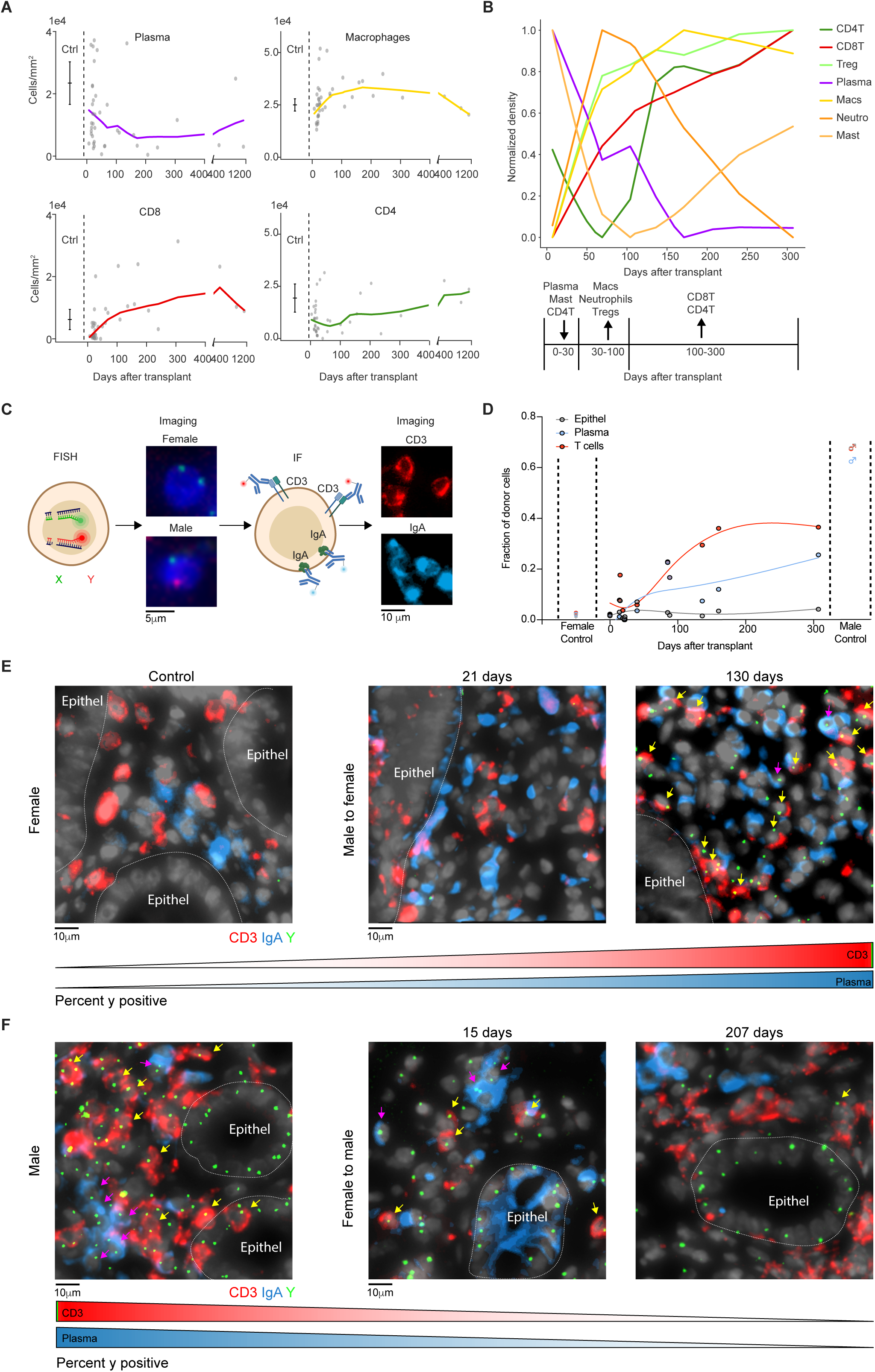
Immune kinetics following transplantation shows persistent host T cells and plasma cells. **(A)** For all patients (dots), shown are the densities of different cell types (y-axis) at the time the biopsy was taken after transplantation (x-axis). Dashed line separates between control and GVHD patients. Solid lines show the lowest fit to the data. **(B)** Lowess-smoothed cell densities as a function of days after transplantation. **(C)** Experimental setup for FISH-IF. Biopsies are first stained with probes for the X and Y chromosomes and imaged. Next, IF is performed for IgA, CD3 and Pan Keratin. FISH and IF images are aligned based on nuclear staining (DAPI). **(D)** Analysis of 10 female patients who received a transplantation from a male donor. For each patient, shown are the fractions of donor cells (male, y-axis) out of the epithelial cells (gray), Plasma cells (blue) or T cells (red) as a function of the number of days between the transplantation and the biopsy (x-axis). Dashed lines separate between control and GVHD patients. Solid lines show the cubic spline fit of the data. **(E)** Representative FISH-IF images of a female control (left) and male donors transplanted into females at either 21 days (middle) or 130 days (right) post transplantation. **(F)** Representative FISH-IF images of a male control (left) and female donors transplanted into males at either 15 days (middle) or 207 days (right) post transplantation. Arrows indicate Y chromosome positive T cells (yellow) and plasma cells (purple).

The kinetics that we observed in the tissue corresponded with previously described kinetics of immune reconstitution by donor cells in the blood ^53^. However, recent studies of GVHD focusing on the recovery of T resident memory (Trm) cells in the skin and the GI tract in non-human primates indicated that host Trm cells can persist in tissues even years post-transplantation, and even if their levels in peripheral blood are below detection level ^17,31,32,34^. As such, we wished to explore in our cohort of GVHD in the gut what fraction of the different immune cells are from host or donor origin, and how these fractions change as a function of time after transplantation.

To distinguish between donor and host-derived cells, we focused on a subset of 10 patients from our cohort in which the host was female and the donor was male (M->F). These patients were sampled at various time points after transplantation, from several weeks to several months, and present a good representation of the entire cohort (**Fig. 1A**). We performed FISH against the X and Y chromosomes to distinguish donor and host cells. We then performed cyclic staining with fluorescently conjugated anti-CD3, anti-IgA, and anti-keratin antibodies to classify T cells, plasma cells, and epithelial cells respectively. We then quantified the percentage of cells that were of male origin (**Fig. 5C**). To assess the sensitivity and robustness of the approach we first stained control samples. We found in male samples that on average 65% of plasma, T cells and epithelial cells were positive for the Y chromosome, while in female samples Y chromosome positive cells were almost undetectable with an average of 0%. This indicates that our assay has 100% specificity and 65% sensitivity (**Fig. 5D, E-F left panel**).

To chart the origin of immune cells in GVHD patients, we evaluated the percentage of T cells (CD3+) and plasma cells (IgA+) in the lamina propria that were of male origin in M->F samples (**Fig. 5D, E**). We found that in the first month following transplantation we could detect ample T cells and plasma cells in the gut, mainly originating from the host (**Fig. 5E, 21 days**). The fraction of donor cells increased in patients with longer times since their transplantation (**Fig. 5E, 130 days**), mirroring the dynamics that we observed in the MIBI-TOF and the dynamics that were reported in blood. Samples in which the host was male, and the donor was female (F->M) showed a similar trend (**Fig. 5F**). Notably, complete replacement of the T cell and plasma cell compartments were not achieved, and we could still detect host T cells and plasma cells even more than 300 days after transplantation (**Fig. 5D)**.

Altogether, our FISH results support the observations derived from MIBI regarding differential dynamics of immune cells in the gut. They suggest that initiation of GVHD in the gut may occur before reconstitution of donor T cells and that host T cells persist in the tissue for a long period of time after transplantation. This supports previous observations ^31,32,34^, suggesting that the underlying mechanisms of GVHD are not limited to donor-derived T cells.

### Inter-crypt heterogeneity associates with changes in the local immune microenvironment

Next, we leveraged our spatial data to explore how immune populations are distributed within the gut. To this end, we used the segmentations of individual crypts and villi, and for each performed a 75-pixel dilation to quantify the prevalence of distinct immune populations in their immediate microenvironment (**Fig. 6A, methods**). As expected, this analysis revealed significant inter-patient heterogeneity in the prevalence of different immune cells, mirroring our results at the patient level. However, we also observed significant intra-patient heterogeneity in the prevalence of different immune cells in the vicinity of individual crypts and villi. The variance within a single patient sometimes spanned the entire variance observed in the cohort (**Fig. 6B, Fig. S4A**). **Figure 6C** shows an example from one patient who had a dense concentration of CD8T cells next to two of the crypts. Interestingly, these crypts are also the most deteriorated, as manifested by their reduced size and altered morphology. Other crypts in the immediate vicinity are intact, with little infiltration of T cells. **Figure 6D** shows a similar trend for neutrophils in a different patient. We observed a high local density of neutrophils in one specific crypt, which was small, isolated and morphologically deteriorated. Other crypts in the immediate surroundings had no infiltration of neutrophils and showed normal morphology. This trend was recapitulated in more patients from the cohort (**Fig. S4B**)

**Figure 6:**
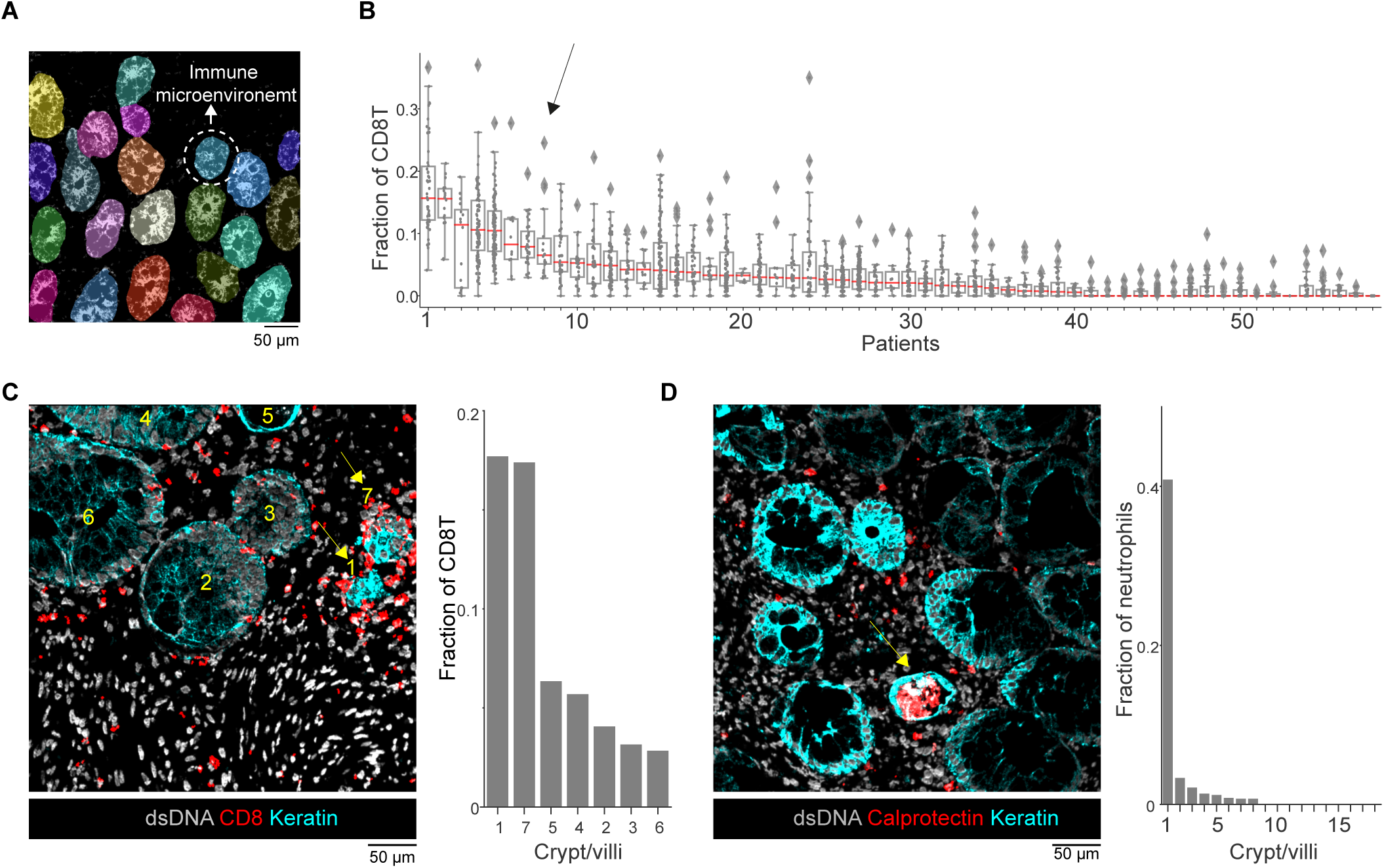
Changes in the local immune microenvironment associate with inter-crypt heterogeneity and GVHD manifestation. **(A)** Individual crypt/villus objects were segmented using Cellpose and dilated to quantify their local immune microenvironments. **(B)** Fraction of CD8T cells out of the immune cells surrounding individual crypts. Box plots summarize all crypts in individual patients and are between the 25th and 75th percentiles. Dots represent individual crypts. Red line indicates the median. Diamonds represent outliers. Arrow indicates a patient with large heterogenity in the fraction of CD8T cells with several outliers. **(C)** Left: example image from the patient marked by an arrow in B, showing the distribution of CD8T cells (red) relative to individual crypts (Keratin, cyan). Seven individual crypts are marked by numbers. Arrows indicate deformed crypts that are enriched with CD8T cells. Right: quantification of the fraction of CD8T cells in the vicinity of the crypts shown in the image. **(D)** Left: example image showing the distribution of neutrophils (calprotectin, red) relative to individual crypts. Arrow indicates a deformed crypt enriched with neutrophils. Right: quantification of the fraction of neutrophils out of all immune cells in the vicinity of the crypts shown in the image.

These results support previous observations that have defined GVHD as a ‘patchy disease’ ^54^. Moreover, they suggest that this patchiness may be driven by local immune microenvironments that are formed during the course of the disease. While it is unclear how these local microenvironments are maintained, the increase in fibrosis and elevation in collagen that we observe in the microenvironment (**Fig. 3F**) could potentially play a role in this process.

### A multiplexed imaging-based predictor for clinical classification of GVHD

We evaluated whether multiplexed-imaging of the immune microenvironment could identify important attributes that determine the severity of the disease. To this end, we combined features extracted from the cohort including cell composition and cell phenotyping and applied principal component analysis (PCA) to summarize the associations between the features and the clinical data. Since the colon has a different immune composition than the duodenum (^40^ and **Fig. 3B**) and in our cohort colon biopsies were associated with increased disease severity, we discarded them from the analysis to avoid confounding factors. The principal components of the data stratified the control group from the mild and severe GVHD **(Fig. 7A)** where the feature that is most associated with PC1 is immune infiltration (variance 17%) and the feature that is most associated with PC2 is the stromal cells (variance 14%) (**Fig. S4C**). We further focused only on the GVHD samples and annotated the patients on a PCA plot according to both the clinical and pathological severity **(Fig. 7B)**. We found that the first PC separated clinically mild and severe patients. Mild patients associated with more epithelial cells, mast cells and CD103+ CD8T cells, while severe patients correlated with more macrophages and GZMB+ CD8T **(Fig 7C, D)**.

**Figure 7:**
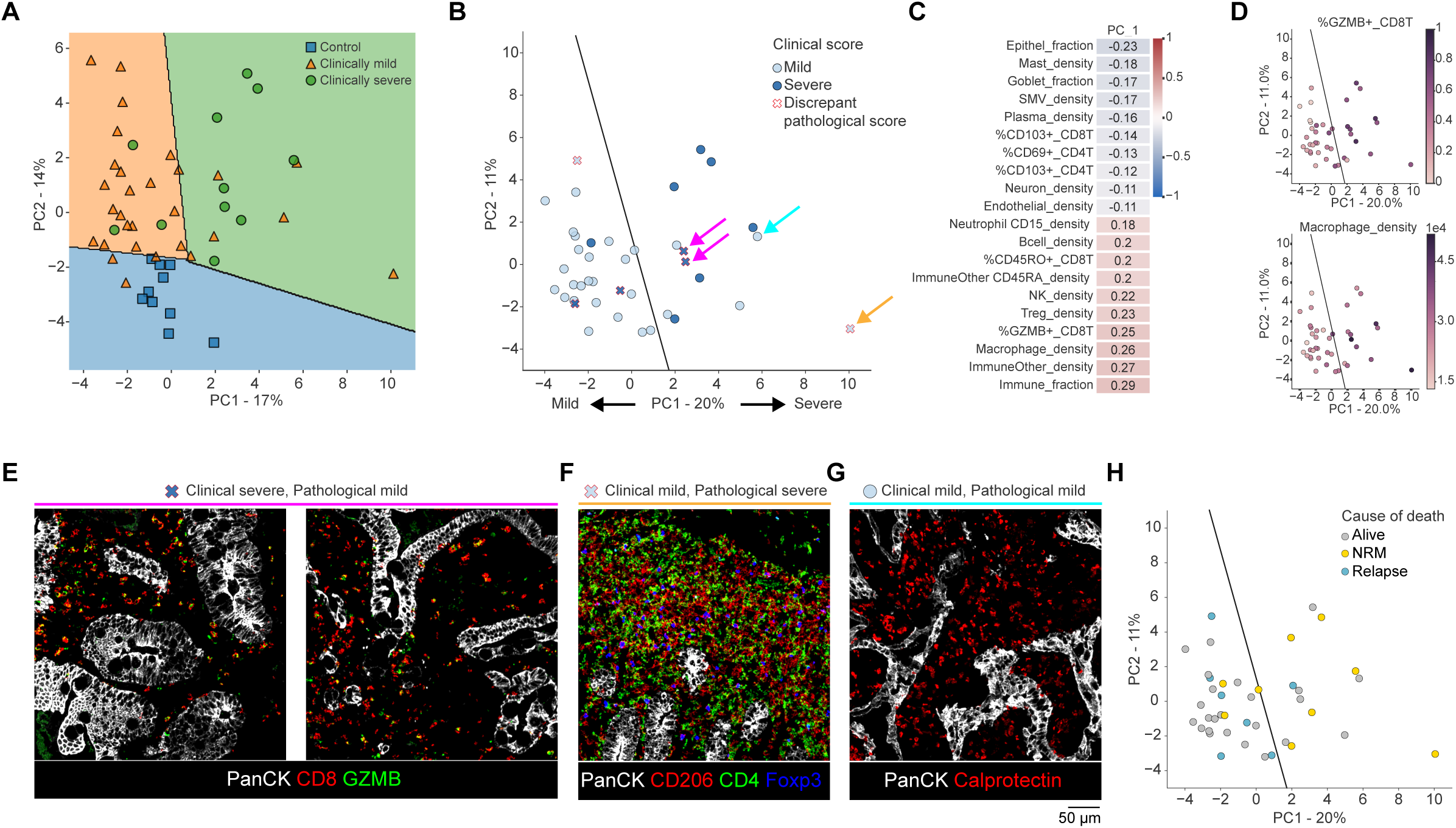
A multiplexed imaging-based predictor for clinical classification of GVHD. **(A)** Principal component analysis (PCA) using all features extracted from the cell composition and phenotype analysis. Axes show the first two PCs and numbers indicate the variance explained. Each dot represents a patient. **(B)** PCA of GVHD patients. The color of the dots indicates the clinical stage of the disease. PC1 is correlated with clinical disease severity. Patients in which the clinical score (severe/mild) and pathological score (severe/mild) disagree are marked by an X. Colored arrows denote patients highlighted in E-G. **(C)** The top ten features contributing to PC-1 in (B). **(D)** PCA of GVHD patients colored by selected features from C. **(E)** Representative images from two patients with severe clinical scores and mild pathological scores (indicated by pink arrows in B). The images demonstrate that although crypt morphology is not deteriorated, there is a high abundance GZMB^+^ CD8T cells. **(F)** The image shows abnormal enrichment of macrophages and Tregs in a patient with mild clinical score but severe pathological score (indicated by an orange arrow in B).**(G)** Enrichment of neutrophils in a patient with both mild clinical and pathological scores, which did not respond to steroids (indicated by a cyan arrow in B). **(H)** PCA of GVHD patients colored by cause of death. Cyan: relapse. Yellow: non-relapse mortality (NRM), which includes GVHD. Gray: patients who survived more than 3 years.

Clinical and pathological diagnosis of GVHD in patients following HSCT is sometimes discrepant ^7,8^. We therefore evaluated how well features extracted from multiplexed imaging agreed with both clinical and pathological scoring. In our cohort, six patients (15%) had a disagreement between the pathological and clinical score (**Fig. 7B**). While this number is too small to perform rigorous statistics, we evaluated the multiplexed images of these patients. Four patients exhibited severe clinical symptoms, but mild pathological scores. Of these, 50% (2 out of 4) showed an enrichment of macrophages and GZMB+ CD8T cells, placing them in the ‘severe’ area of the PCA (**Fig. 7B, pink arrows, Fig. 7E**). An additional discrepancy explained by the PCA involves a patient who was pathologically severe but exhibited only mild clinical symptoms at the time of the biopsy. The patient’s immune microenvironment was abnormally enriched with macrophages and Tregs, and a follow-up revealed that he died from GVHD (**Fig. 7B orange arrow, Fig. 7F**). However, it is important to note that the features collected in this multiplexed imaging study do not explain all mismatches. For instance, two patients who had severe clinical symptoms and mild pathological scores, also seemed mild in our analysis. It is possible that in such patients the biopsy did not capture the inflammatory region, or additional features are necessary to implicate the tissue as diseased.

In addition to the potential of this approach to explain mismatches between pathological and clinical scores, it can also identify patients who might be at risk for severe GVHD, despite lacking any severe pathological or clinical indications. We found four such patients who exhibited features usually observed in severe cases. Of these patients, 50% (2 out of 4) were either non-responders to corticosteroids or non-survivors. For example, one patient who had a major enrichment of neutrophils in the biopsy was also not responsive to corticosteroids (**Fig. 7B cyan arrow, Fig 7G**).

Finally, we examined the cause of death across patients. We divided the patients to those that were still alive three years following transplantation, those that passed from cancer relapse, and those that passed from non-relapse causes, including GVHD and infection. We found that the first PC of our data also correlated with cause of death (Wilcoxon P=0.02, **Fig. 7H**). Patients that passed from non-relapse causes tended to have a more activated immune response, with higher fractions of macrophages, GZMB+ CD8T cells and neutrophils, whereas patients that experienced cancer relapse tended to have less immune allogeneic reactivity activation, presumably leading to a milder graft versus tumor effect (GVT), which could facilitate relapse ^55^.

Altogether, although our cohort lacks statistical power, anecdotal observations indicate that features derived from the multiplexed images may enhance our understanding of patient stratification across various clinical parameters.

## Discussion

GVHD is the most common immune complication of aHSCT, primarily manifesting in the skin, liver and GI tract ^14,24,56^. Here, we used multiplexed imaging to comprehensively characterize the spatial organization of epithelial, stromal and immune cells in 59 diagnostic biopsies of patients with gastrointestinal GVHD and 10 healthy controls. We found that normal duodenum displayed various levels of structure and was highly stereotypic across individuals. Recent studies showed that the intestinal epithelium is functionally organized, with gene expression gradients both along and perpendicular to the crypt-villi axis ^41,42^. Comparing transcriptomics and proteomics in mouse duodenum showed that the zonation patterns of mRNAs and proteins along the epithelium were often not correlated ^42^. Here, we identified two epithelial proteins that were zonated along the crypt-villus axis (HLA-II) and baso-apical axis (Na^+^/K^+^-ATPase) across individuals. Moreover, we uncovered zonation of immune cells in the gut, in which CD4T cells, plasma cells and macrophages were spatially organized from the crypts to the tip of the villus. This recurrent organization between individuals raises questions regarding the mechanisms that drive it, as well as its functional implications in healthy gut homeostasis.

In contrast to healthy duodenum, GVHD showed a heterogeneous morphology, including increased fibrosis, alterations in crypt size and spacing, loss of Paneth cells, and accumulation of endocrine cells in the crypts. Although immune-mediated fibrosis is a hallmark feature of chronic GVHD ^14,24^, we are not aware that its prevalence in acute GVHD has been extensively explored in previous literature. The loss of Paneth cells has been previously described and supports clinical observations showing an increase in the anti-microbial protein REG3A in the plasma of GVHD patients, presumably due to the destruction of Paneth cells ^50,51^. This reduction in Paneth cells was accompanied by an increase in endocrine cells, specifically near the muscularis mucosa, where Paneth cells normally reside ^10–13^. Similar increases in endocrine cells have been reported in Crohn’s disease and ulcerative colitis ^57,58^. These results raise several hypotheses. The first is that Paneth cells are preferentially attacked in GVHD, at least in a subset of patients. This hypothesis corresponds with recent observation in mice, which suggested that the intestinal stem cells are the primary target of attack in GVHD ^9,59^. In humans, either Paneth cells are preferentially attacked, or they could be secondary targets due to their spatial proximity to the intestinal stem cells. Since the accumulation of endocrine cells was observed primarily near the base of the crypt, another intriguing possibility is that GVHD causes misdifferentiation of intestinal stem cells to endocrine cells instead of Paneth cells. Differentiation to distinct types of epithelial cells is largely controlled by complex interactions between wnt and notch-delta signaling ^60^. Misregulation in these pathways could drive an accumulation of endocrine cells. Additional spatially resolved transcriptomic and proteomic studies to examine these pathways could shed light on the mechanisms driving epithelial deformation in GVHD, as well as on the subtypes of endocrine cells that are increasing ^61,62^.

The pathophysiology of GVHD typically implicates an initiation phase, in which damage from the conditioning regime triggers the release of damage-associated molecular patterns (DAMPs) and pathogen-associated molecular patterns (PAMPs); a T cell activation phase, in which host APCs activate alloreactive donor T cells; and an effector phase, in which effector T cells and pro-inflammatory cytokines damage the epithelial cells of the gastrointestinal (GI) tract ^14,63^. However, several recent studies have begun to challenge this simplified paradigm, implicating a role for plasma cells ^19,64–66^, macrophages ^64,67,68^ and neutrophils ^28,69–71^ in the initiation and progression of the disease. While most works focused on specific cell types, here we took a holistic approach and found evidence to support the involvement of several types of immune cells, in addition to T cells, in disease initiation and progression. Patients with clinically severe GVHD generally had less plasma cells and CD4T cells, and an increased number of macrophages and neutrophils. Decreased plasma cells in GI-GVHD have also been recently recognized by Jarosch and colleagues ^19^. Interestingly, when analyzing patients according to the time after transplantation, we identified stereotypical dynamics, with a rapid drop in plasma cells immediately after the transplantation, a slower increase in macrophages, Neutrophils and T regulatory cells, and finally an accumulation of effector T cells. These dynamics were reminiscent of previously described differential reconstitution dynamics in the blood ^53^. They suggest that host plasma cells, important for maintaining the immune barrier in the gut, are rapidly reduced following the conditioning regimen, but slowest to reconstitute from the donor hematopoietic stem cells. This shift from homeostasis may increase epithelial vulnerability and serve as another driver of the disease. Further mechanistic exploration is necessary to establish the importance of the role that plasma cell depletion may play in disease initiation.

To follow the dynamics of donor immune cell reconstitution in the gut we performed FISH of the X and Y chromosomes to distinguish between donor and host immune cells in sex-mismatched samples. This experiment yielded several intriguing results. We found that unlike reports on the blood, in which the host immune system is almost completely ablated by the conditioning regimen ^53^, in the gut host cells dominated the plasma and T cell compartments for extended periods of time following transplantation. In some of the patients diagnosed with GVHD shortly after transplantation, we could hardly identify any host T cells in the gut, suggesting that they may not be the sole driver of the disease in these patients. Accordingly, in our cohort, host T cells and plasma cells persisted in the gut even several months after transplantation. Finally, the reconstitution dynamics were different for the T cells and the plasma cells, matching their differential reconstitution dynamics in the blood ^53^. These results mirror recent observations in the skin and gut, which have reported similar findings for T cells and suggested that donor antigen-presenting cells may activate host T cells to instigate a local inflammation ^31,32,34^.

Pathologically, GVHD sometimes manifests in ‘patchy’ or ‘diffuse’ patterns ^72^. In our analyses, we also identified inter-crypt heterogeneity in disease progression, with deteriorated crypts located next to crypts with normal morphological appearance. Interestingly, we also saw that some of these crypts were characterized by a unique local immune microenvironment, which was enriched for either activated CD8T cells, Tregs, Neutrophils or macrophages. This immune concentration suggests that many cell types could be participating in either driving the disease or amplifying local damages. Moreover, the localized nature of these immune microenvironments and the proximity of destructed and healthy crypts cannot be explained solely by the classic view of donor T cell activation by host APCs. It implicates the existence of local processes, barriers and feedback loops that drive this compartmentalization. Our analysis suggests that one factor that contributes to this compartmentalization could be fibroblasts that secrete ECM proteins, which limit the motility of immune cells in the tissue. Additional mechanisms which require further investigation include a local breach of the mucosal barrier, which facilitates local entry of bacteria and local immune activation ^23^.

One of the long-standing clinical dilemmas in treatment of GVHD is that pathological grades are only moderately correlated with the clinical grade, which is based on symptomatic disease severity ^7,8^. In our cohort, six patients had discrepant clinical and pathological scores. In three of these (50%), we identified immune features that are usually observed in severe patients, including the presence of GZMB+ CD8T cells and high abundance of macrophages. We hypothesize that in these patients we can already identify the immune hallmarks driving the disease, but epithelial deterioration was either not yet established, or missed by the biopsy due to patchy appearance of the disease. It is thus possible that augmenting H&E staining with immunohistochemistry for a select number of proteins, either multiplexed or in isolation, could serve for more predictive and sensitive pathological diagnosis. In the future, it may be further possible to tailor therapies to the specific immune composition in each patient. To explore such exciting possibilities, larger cohorts need to be profiled. Altogether, this study provides a comprehensive spatial characterization of the immune microenvironment of GVHD in the GI and lays the foundation for additional work to follow in GVHD, inflammatory bowel diseases and cancer.

## Methods

### Samples and study participant details

GVHD samples: Gut biopsies (59) were obtained from 57 patients during routine diagnosis of acute GVHD at Hospital St Louis, Paris and have not been collected specifically for this study. Two patients had two biopsies taken at different times and were treated independently for the purpose of this study. All patients have given consent to allow research on these samples. The study was approved by IRB 00003888; project number 21-799. Weizmann IRB 1189-1 and and 2318-1. All patients’ pathology reports were supervised by PB (Prof of Pathology; University of Paris, France). Control Duodenum samples were obtained from 10 patients that underwent Whipple procedure, whereby a tumor from the head of the pancreas is removed together with healthy duodenum under MTA FO-TB-1979. FFPE tissue blocks were retrieved from the tissue archive at the Institute of Tissue Medicine and Pathology (ITMP), University of Bern, Switzerland. Normal GI tissue regions were annotated on corresponding hematoxylin and eosin (H&E)-stained sections by a board-certified surgical pathologist (M.W.). A next-generation TMA with 2 mm diameter cores was assembled using a TMA Grand Master automated tissue microarrayer (3DHistech). In addition, 4 duodenum samples were collected directly in the operations of patients undergoing Whipple procedure at Sheba Medical Center, Israel. Use of tissue was approved by Sheba Medical Center Helsinki committee (approval number 866521SMC). Samples were taken only in cases in which no pathology was observed in the small intestine. ∼5 cm intestinal tubes from the ligament of Treitz were resected and longitudinally opened to expose the mucosa surface. Tissues were gently washed in PBS and mucosal sheets were sectioned by performing mucosectomy. Sheets were rolled to a Swiss roll to maximize tissue space. Full thickness intestinal layers were fixed in pre-chilled 4% FA for 24 hours followed by fixation in 1% PFA for preparation of FFPE blocks.

### MIBI-TOF staining

Tissue sections (5 µm thick) were cut from FFPE tissue blocks of the biopsies using a microtome, and mounted on gold-coated slides (Ionpath, Menlo Park, CA) for MIBI-TOF analysis. Slide tissue sections were baked at 70°C for 20 min. Tissue sections were deparaffinized with 3 washes of fresh xylene. Tissue sections were then rehydrated with successive washes of ethanol 100% (2x), 95% (2x), 80% (1x), 70% (1x), and ultra-pure DNase/RNase-Free water (Bio-Lab, Jerusalem, Israel), (3x). Washes were performed using a Leica ST4020 Linear Stainer (Leica Bio-systems, Wetzlar, Germany) programmed to 3 dips per wash for 30 s each. The sections were then immersed in epitope retrieval buffer (Target Retrieval Solution, pH 9, DAKO Agilent, Santa Clara, CA) and incubated at 97°C for 40 min and cooled down to 65°C using Lab vision PT module (Thermo Fisher Scientific, Waltham, MA).After cooling to room temperature, slides were assembled with a coverplate on a sequenza rack and were washed 2 times with 1 ml of TBS-T (Ionpath) diluted in ultrapure water. Sections were then blocked for 1h with 300 μl of 5% (v/v) donkey serum (Sigma-Aldrich, St Louis, MO) diluted in TBS-T. Metal-conjugated antibody mix was prepared in 5% (v/v) donkey serum TBS-T wash buffer and filtered using centrifugal filter, 0.1 µm PVDF membrane (Ultrafree-Mc, Merck Millipore, Tullagreen Carrigtowhill, Ireland). Two panels of antibody mixes were prepared. The first panel contained most of the metal-conjugated antibodies and was incubated overnight at 4°C in a humid chamber. The second mix contained antibodies for dsDNA and α-SMA. Following overnight incubation, slides were washed 2 times with 1ml TBS-T and once with 300 μl 5% (v/v) donkey serum (Sigma-Aldrich, St Louis, MO) diluted in TBS-T and incubated with the second antibody mix for 1h at room temperature. Slides were washed 3 times with TBS-T and then were removed from the sequenza assembly. Slides were fixed for 5 min in diluted glutaraldehyde solution 2% (Electron Microscopy Sciences, Hatfield, PA) in PBS-low barium. Tissue sections were dehydrated with successive washes of Tris 0.1 M (pH 8.5), (3x), Ultra-pure DNase/RNase-Free water (2x), and ethanol 70% (1x), 80% (1x), 95% (2x), 100% (2x). Slides were immediately dried in a vacuum chamber for at least 1 h prior to imaging.

### MIBI-TOF imaging

Imaging was performed using a MIBIScope mass spectrometer (Ionpath, Menlo Park, CA) with a Xenon ion source. FOVs of 400µm × 400µm or 800µm × 800µm were acquired using a grid of 1024X1024 or 2048X2048 pixels respectively, with 2 msec or 1 msec dwell time per pixel respectively.

### Dual FISH and immunofluorescence staining

Tissue sections (5 µm thick) were cut from FFPE tissue blocks of the biopsies using a microtome and mounted on slides. Slide tissue sections were baked at 70°C for 1h. Tissue sections were deparaffinized with 3 washes of fresh xylene. Tissue sections were then rehydrated with successive washes of ethanol 100% (2x), 95% (2x), 80% (1x), 70% (1x), and ultra-pure DNase/RNase-Free water (3x). Washes were performed using a Leica ST4020 Linear Stainer (Leica Bio-systems, Wetzlar, Germany) programmed to 3 dips per wash for 30 s each. The sections were then immersed in epitope retrieval buffer (Target Retrieval Solution, pH 9, DAKO Agilent, Santa Clara, CA) and incubated at 97°C for 40 min and cooled down to 65°C using Lab vision PT module (Thermo Fisher Scientific, Waltham, MA). Slides were washed twice for 5 min with PBS (Thermo Fisher Scientific, AM9625) diluted in ultra-pure DNase/RNase-Free water. Sections were marked by a hydrophobic pen (Bar Naor, Israel), then washed in PBS-T (Abcam, AB6247) twice and 2XSSC once, both diluted in ultra-pure DNase/RNase-Free water. Tissues were permeabilized with 10 mg/ml proteinase K diluted in 2XSSC for 10 min in 50°C. Slides were washed in 2X SSC, 5min and dehydrated with successive washes of 70%, 80%,100% ethanol for 2 minutes. Probes for X (KromaTID, USA, Subtelomere CHR Xp and Xq 550 and Subcentromer CHR Xp 550) and Y chromosome (KromaTID, USA, Subtelomere CHR Yp 643) were added on the tissues. Samples were denatured by heating the slide on a hot plate at 75°C, 5min. Slides were incubated overnight with probes in a humid chamber at 37°C. The following day slides were washed with 0.4XSSC 0.05% tween-20 at 60°C for 2 min. Then the slides were washed twice with 2XSSC and once with pure water. To stain the nuclei of the cells, tissues were incubated for 5 min with 150 ng/ml DAPI (Biolegend, 422801) for 5 min at room temp (rt). Slides were washed twice with 2XSSC, mounted with Fluoromount G and closed with a coverslip. Images of sections were acqiured by a widefield microscope (Leica DMI8) with 20X/ 0.8 dry objective. Following imaging, coverslips were incubated in PBS and removed. The tissues were then washed twice in PBS-T. Some of the tissues with high autofluorescence were incubated with TrueView (Vector Laboratories, SP-8400) for 3 min. Following incubation, slides were washed 3 times with PBS-T and then blocked with blocking buffer (PBS-T 5% NDS) for 1h at rt. Sections were stained overnight with 1.25 μg/ml anti-CD3 alexa 647 (CST, CST-63178S). The following day slides were washed with PBS-T and incubated for 1h at rt with 15 μg/ml fab fragment anti-rabbit alexa 647 (Jackson ImmunoResearch, 711-607-003). Slides were stained overnight with 33 μg/ml anti-IgA alexa488 (Abcam, ab223410) and 5 μg/ml anti-keratin alexa 550 (Abcam, ab279324). The following day slides were washed twice with PBS-T and stained with 150 ng/ml DAPI for 5 min at rt. Slides were washed twice with PBS-T. Slides were mounted with Fluoromount G and closed with a coverslip. Images of sections were acqiured once again by a widefield microscope (Leica DMI8) with 20X/ 0.8 dry objective. Images were aligned with a correlation algorithm based on DAPI staining (Matlab). Cells were segmented with cellpose 2 plugin in QuPath^73^. Cells were classified by merging 3 single channel classifiers in Qupath. Cell classifications were then manualy inspected and corrected when needed. X and Y spots were counted using QuPath with the subcellular detection mode.

### Antibodies

A summary of antibodies, reporter isotopes, and concentrations can be found in Supplementary Table 2. Conjugated primary antibodies were purchased from Ionpath or conjugated in-house using the Maxpar X8 Antibody Labeling Kit (Fluidigm, California, USA) or MIBItag Conjugation Kit (Ionpath) according to the manufacturer’s recommended protocols.

### Low-level analysis

Multiplexed images were processed to improve image quality including background subtractions, noise removal, and aggregate filtering. Removal of channel background was done using a custom in-house script that takes two channels, a source and a target, and iteratively subtracts the scaled source image from the target with the goal of maximizing the mean squared error between them which is equivalent to the largest distance between the images. The resulting images were manually inspected to ensure that the approach is sufficient to remove background related noise while keeping the real signal. Noise removal and aggregate filtering were performed using MAUI pipeline as previously described^74^.

### Cell segmentation

Cell segmentation was performed using Mesmer^75^ which is wrapped within the ark-analysis toolbox (https://github.com/angelolab/ark-analysis) with the adaptation for segmenting specific cell types. dsDNA was used as a nuclear channel and Pan-keratin, E-cadherin, Na^+^/K^+^-ATPase, CD45 and HLA-class-I as membranal channels to generate whole cell segmentation. SMA and CD68 were also added as nuclear channels to capture muscle and macrophages in which the nuclei are often not visible in the plane of imaging. Since goblet cells are larger than most cells in the GI and have a unique structure, they were segmented separately. To this end, Mesmer was run again with Mucin as the nuclear channel and epithelial channels for membrane detection. The rescale factor of Mesmer was adjusted to 0.4 to down-scale the goblet cells to the range of sizes that Mesmer was trained on. Finally, we combined the segmentation output from the two iterations to create cell segmentation masks for each image.

### Generation of region masks

Region masks were generated using a pixel classifier in ilastik^76^ to annotate anatomical structures in the intestine including, the crypt-villi region, muscle region, smooth muscle vasculature (SMV) and Brunner glands. For each region, the appropriate channels were used as features for the classifier, which was trained on user provided annotations. Crypt-villi region: max(Keratin, Ecad, Na^+^/K^+^-ATPase) + CD45; muscle region: SMA + Collagen + max(Vimentin, Keratin, Ecad, Na^+^/K^+^-ATPase); SMV: SMA + CD31 + Epithelial channels; Brunner glands: Mucin + max(Keratin, Ecad, Na^+^/K^+^-ATPase) + Lysozyme. In addition, regions of the submucosa and follicles were annotated manually in QuPath due to their sparsity in the images. Finally, a mask of the lamina propria region was generated by taking the negative of the union of all other region masks.

### Cell classification

Cell classification was performed using a custom active learning pipeline developed in-house. Spatial expression features were extracted from the images for each cell and joined in a cell table. These features included the mean intensity of proteins as well as the counts of non-zero pixels within the cell segmentation, within dilated (10 pixels, 50 pixels) and eroded (5 pixels) cell segmentations, and within neighboring cells. Region masks were also added as additional features to the cell table. Cell types were determined manually by inspecting the mean protein expression of clusters generated with flowSOM^77^. Initial labels were assigned to a subset of cells using stringent gating criteria based on protein expression levels. Two gradient-boosted decision tree models^78,79^ were then trained on the labeled data. Cells with differing predictions from these models were sampled for further manual annotation, to focus on areas of uncertainty to improve the model. Newly labeled cells were added to the training data, and the process was iteratively repeated 17 times to progressively refine the model’s performance in cell classification. In total 14k cells were labeled including the initially gated cells and additional manually labeled cells. The final classification was reviewed by manual inspection of cells from 20 crops as well as by comparing the protein expression in full FOVs to colored cell type maps. Segmented goblet cells located in the lumen were filtered out from the cell table as they may represent secreted mucin rather than actual cells. Also, goblet cells with area smaller than 60 pixels (9μm^2^) were removed.

### Cell composition and phenotyping

The relative cell abundances were calculated out of the total cells per FOV, or from of the major lineages, including immune, epithelial and stromal cells. The density in the lamina propria was calculated by counting the cells in the lamina propria region mask, normalizing by the area of the mask and converting the value to cells per mm^2^. Similarly, the density in epithelium was normalized by the area of the epithelium region mask. In each analysis the appropriate measurement was used as indicated in the figure legends. For phenotyping analysis, the expression table was generated as follows: for each cell and each protein, the summed intensity was computed and normalized by the cell area and then the values were arcsinh-transformed. Cells were noted as positive if this value was greater than 1. Finally, the relative abundance of the positive cells was calculated for each corresponding cell type. Patients with less than 20 cells per cell type were removed from the analysis to reduce noisy values.

### Zonation of protein expression and cell types

Selected control images were computationally rotated to orient the villi vertically from base to tip. The images were then divided into 20 vertical bins spanning from the base to the tip of the crypt-villi mask. For each bin, the percentage of positive pixels for E-cadherin HLA-I, and HLA-II was calculated relative to the total pixels within the mask. For immune zonation, 7 FOVs from 4 patients were included due to their structure capturing the entire villi from base to tip. Images were rotated and the vertical (y) distance of the centroid of the cells was used to determine the distance of the cell along the vertical villi axis. To aggregate cells from different FOVs, the y distance for each fov was normalized to the approximate length of the villi using a min-max normalization based on the mean y distance of Paneth cells at the base and the 99.9^th^ percentile of cell distances.

### Endocrine distance to muscle analysis

For this analysis endocrine and neuroendocrine cells were aggregated and will be henceforth referred to as endocrine cells. The mean distance of endocrine centroids to the muscle mask was calculated and tested against the baseline distance of epithelial cells from the muscle. To generate the baseline distance, the locations of endocrine cells were permuted within epithelial cells in the image and the average distance to the muscle was computed. This process was repeated 500 times to generate a null distribution, and the z-score was calculated to test the deviation of the real endocrine distance from the null distribution. Images with muscle masks containing less than 50 pixels were excluded from the analysis as well as images with less than 100 epithelial cells to avoid over sampling. Overall, 92 FOVs from control patients (*n*=8) and duodenum GVHD patients (*n*=37) were included in analysis.

### Single-Crypt/villi segmentation

Cellpose2 extension within QuPath was used to segment individual crypt and villi objects in the image. Input images for Cellpose were generated by summing the images of keratin and mucin. To achieve accurate segmentation, parameters of object size and subsampling were fine-tuned. The resulting segmentations were manually curated and any unsegmented crypts/villi were manually labeled using the QuPath annotation tool. For each object, morphological features were calculated, including area, perimeter, circularity, and isolation score. The isolation score is the percentage of the object’s non-overlapping area with neighboring crypts within a 75-pixel expansion ring. To ensure that individual crypts are analyzed, objects with a low circularity score and large perimeter were filtered out.

### Crypt heterogeneity analysis

To investigate the local enrichment of immune cells, a local environment region mask was generated for each object. For CD8T cells, an inner ring of 25 pixels (±10 μm) and an outer ring of 75 pixels (± 30 μm) were combined to form a single region mask, accounting for cells in the lamina propria and IELs. This mask was then intersected with the LP region mask to include only cells within the LP, excluding cells from neighboring crypts. For neutrophils, the entire space inside the object and a 25-pixel outer ring were included, as neutrophils were observed occupying the inner space of the crypts. The relative abundance of each cell type was computed based on the total cells in the environment regions.

## Data availability

All images and annotated single-cell data are deposited in Mendeley’s data repository and can be accessed via this link: https://doi.org/10.17632/j4bscsgn6x.1.

## Acknowledgements

The authors thank Dr. Yoseph Addadi, Dr. Ofra Golani and Dr. Oshrat Galibov Levi for their assistance with fluorescent imaging and analysis, Calanit Raanan, Marina Cohen and Lena Prichislov for histology services. The authors acknowledge the Translational Research Unit at the Institute of Tissue Medicine and Pathology, University of Bern, Switzerland, for providing human tissue samples and excellent technical support with the construction of the TMA. I.M. was supported by a EU - Horizon 2020 - MSCA Individual Fellowship (890733). L.K. holds the Fred and Andrea Fallek President’s Development Chair. She is supported by the Enoch foundation research fund, the Abisch-Frenkel foundation, the Rising Tide foundation, the Sharon Levine Foundation and grants funded by the Schwartz/Reisman Collaborative Science Program, European Research Council (948811), the Israel Science Foundation (2481/20, 3830/21) within the Israel Precision Medicine Partnership program.

COI :

D.M. received research grant from Novartis, CSL Behring and Sanofi, and consulting fee for Novartis, Incyte, Jazz Pharmaceutical, Therakos and Sanofi. G.S. is the recipient of an unrestricted research grant from Alexion.

