## Supplementary Figures for "A spatial atlas of human gastro-intestinal acute GVHD reveals epithelial and immune dynamics underlying disease pathophysiology"

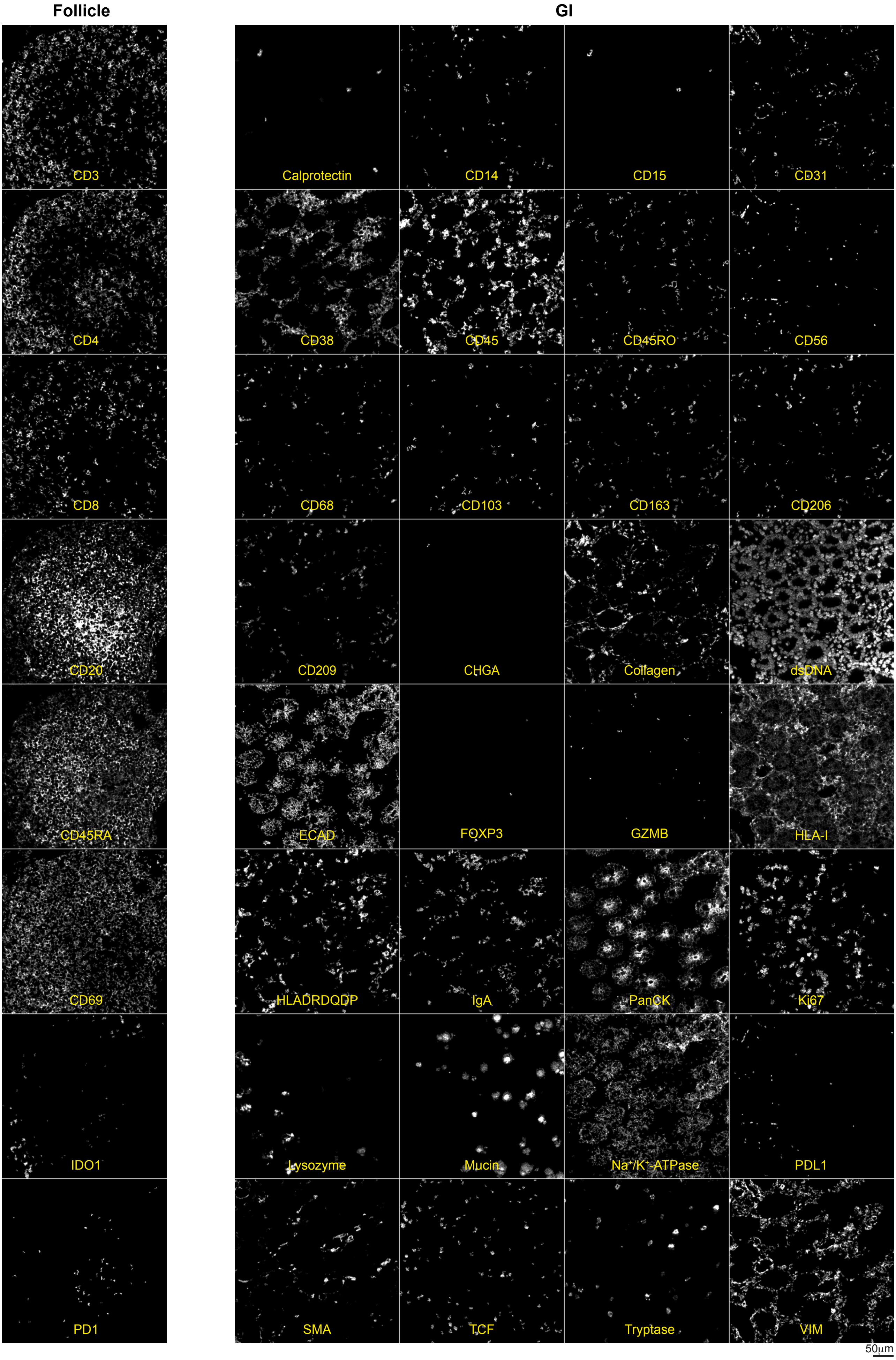


**Figure S1. Calibration of a 40-plex antibody panel to profile human duodenum**

Antibody calibration was performed on control tissues with known staining patterns. Shown are representative MIBI-TOF images of normal human duodenum stained with the antibody panel, depicting an isolated lymphoid follicle (left) and a region from the mucosa (right).

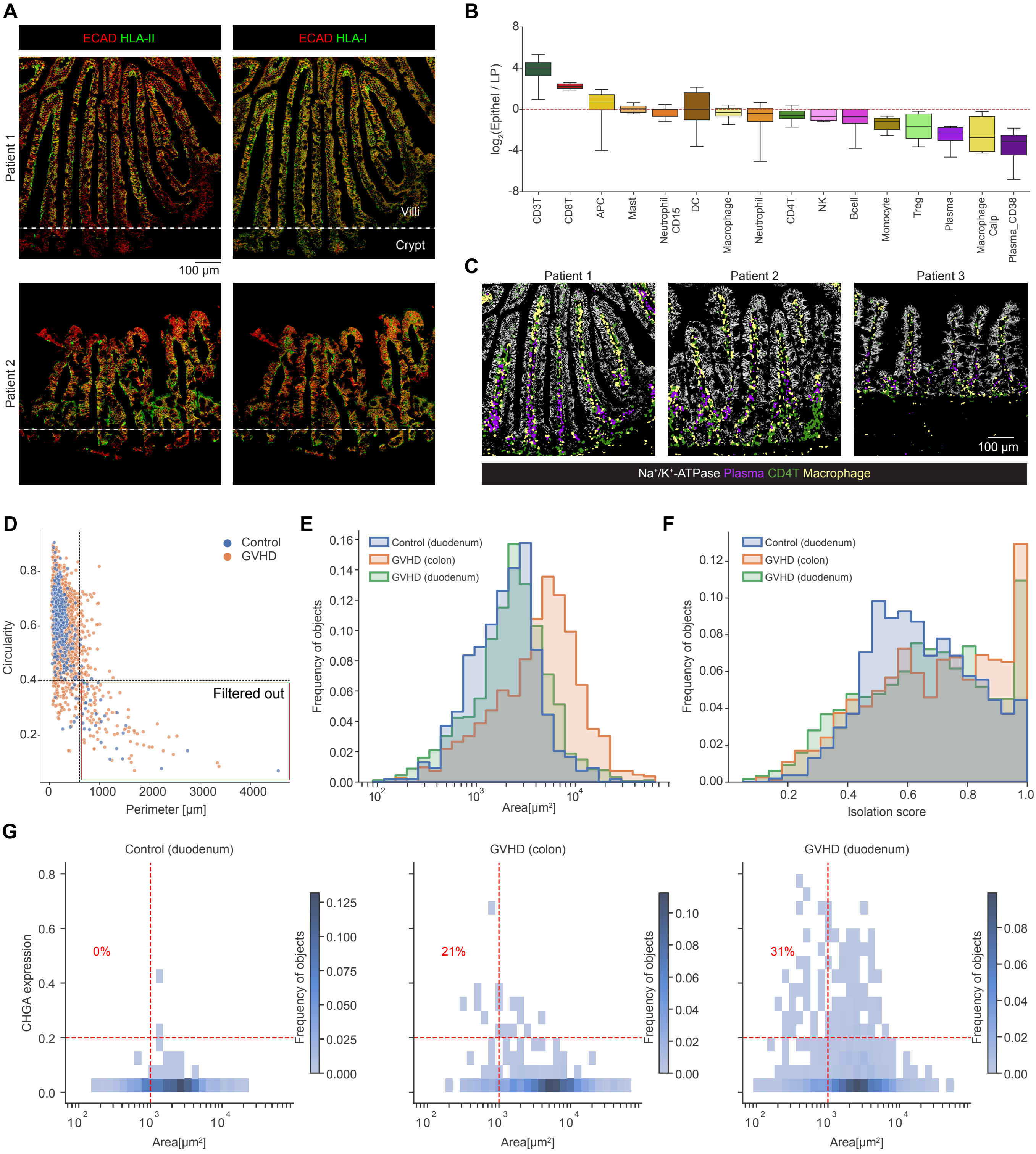

**Figure S2. Spatial immune and epithelial organization in healthy and diseased intestine**

**(A)** Images of healthy duodenum from two individuals (rows) show the expression of E-cadherin (ECAD), HLA-I (left), and HLA-II (right) along the crypt-villi axis. Dashed lines separate the crypts (bottom) and villi (top). **(B)** For each immune cell type (x-axis), shown is a boxplot of the log2-ratio between the fraction of that cell type out of all immune cells inside the epithelium versus the lamina propria (y-axis) across 10 control patients. The dashed red line indicates equal fractions. CD3T and CD8T cells are enriched inside the epithelium. **(C)** Immune zonation of CD4T cells, plasma cells and macrophages along the crypt-villi axis in three control patients. **(D)** A scatter plot showing the perimeter (x-axis) versus circularity (y-axis) of all segmented crypt/villi objects. Each dot represents an object and is colored by the patient (control/GVHD). Objects at the bottom-right quadrant are large with non-circular shapes, which mostly include two or more epithelial objects that were segmented together and therefore filtered from further object-related analyses. **(E)** Histograms of the distribution of crypt/villi sizes in control (blue), colonic GVHD (orange), and duodenal GVHD (green). **(F)** For each crypt/villus object, an isolation score was computed by calculating the percentage of the object's non-overlapping area with neighboring crypts within a 75-pixel expansion ring. Shown are histograms of the isolation score of crypt/villi objects in control (blue), colonic GVHD (orange), and duodenal GVHD (green). **(G)** 2D histograms displaying the sizes of crypt/villi objects (x-axis) and the expression of chromogranin A (CHGA) relative to keratin expression in the objects (y-axis). Bins are colored by prevalence. Dashed red lines highlight small (shrunk) objects with a high expression of CHGA found at the top-left quadrant. The numbers indicates the percentage of patients with at least one object in that quadrant out of all patients. The histograms are separated to control (left), duodenal GVHD (middle) and colonic GVHD (right).

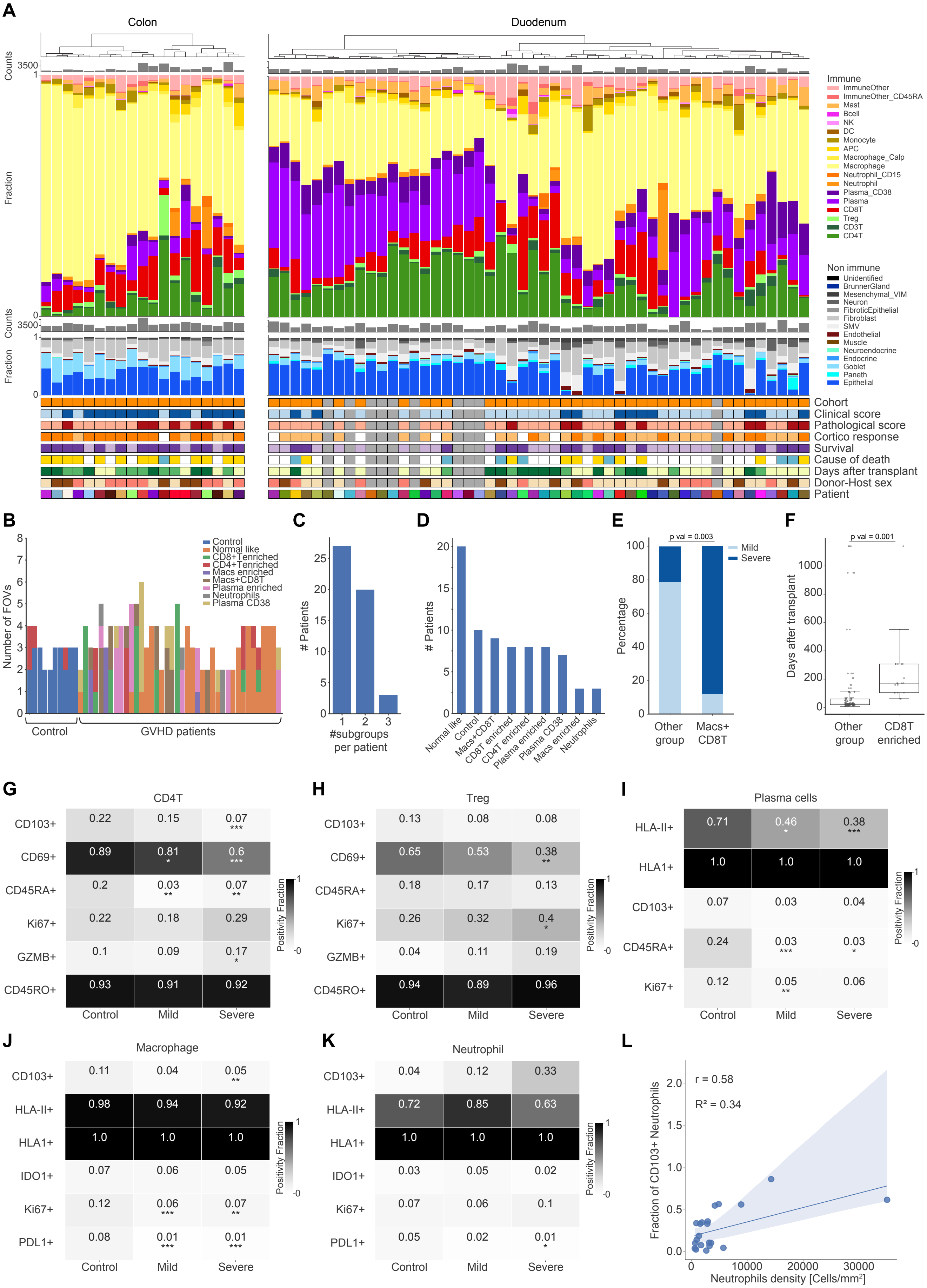

A
Colon
Duodenum
Counts
3500
Fraction
Immune
ImmuneOther
ImmuneOther_CD45RA
Mast
Bcell
NK
DC
Monocyte
APC
Macrophage_Calp
Macrophage
Neutrophil_CD15
Neutrophil
Plasma_CD38
Plasma
CD8T
Treg
CD3T
CD4T
Non immune
Unidentified
BrunnerGland
Mesenchymal_VIM
Neuron
FibroticEpithelial
Fibroblast
SMV
Endothelial
Muscle
Neuroendocrine
Endocrine
Goblet
Paneth
Epithelial
Cohort
Clinical score
Pathological score
Cortico response
Survival
Cause of death
Days after transplant
Donor-Host sex
Patient
B
Number of FOVs
Control
Normal like
CD8+Tenriched
CD4+Tenriched
Macs enriched
Macs+CD8T
Plasma enriched
Neutrophils
Plasma CD38
Control
GVHD patients
C
### Patients
#subgroups per patient
D
### Patients
Normal like
Control
Macs+CD8T
CD8T enriched
CD4T enriched
Plasma enriched
Plasma CD38
Macs enriched
Neutrophils
E
p val = 0.003
Percentage
Mild
Severe
Other group
Macs+ CD8T
F
p val = 0.001
Days after transplant
Other group
CD8T enriched
G
CD4T
CD103+ 0.22 0.15 0.07 ***
CD69+ 0.89 0.81 * 0.6 ***
CD45RA+ 0.2 0.03 ** 0.07 **
Ki67+ 0.22 0.18 0.29
GZMB+ 0.1 0.09 0.17 *
CD45RO+ 0.93 0.91 0.92
Control Mild Severe
Positivity Fraction
H
Treg
CD103+ 0.13 0.08 0.08
CD69+ 0.65 0.53 0.38 **
CD45RA+ 0.18 0.17 0.13
Ki67+ 0.26 0.32 0.4 *
GZMB+ 0.04 0.11 0.19
CD45RO+ 0.94 0.89 0.96
Control Mild Severe
Positivity Fraction
I
Plasma cells
HLA-II+ 0.71 0.46 * 0.38 ***
HLA1+ 1.0 1.0 1.0
CD103+ 0.07 0.03 0.04
CD45RA+ 0.24 0.03 *** 0.03 *
Ki67+ 0.12 0.05 ** 0.06
Control Mild Severe
Positivity Fraction
J
Macrophage
CD103+ 0.11 0.04 0.05 **
HLA-II+ 0.98 0.94 0.92
HLA1+ 1.0 1.0 1.0
IDO1+ 0.07 0.06 0.05
Ki67+ 0.12 0.06 *** 0.07 **
PDL1+ 0.08 0.01 *** 0.01 ***
Control Mild Severe
Positivity Fraction
K
Neutrophil
CD103+ 0.04 0.12 0.33
HLA-II+ 0.72 0.85 0.63
HLA1+ 1.0 1.0 1.0
IDO1+ 0.03 0.05 0.02
Ki67+ 0.07 0.06 0.1
PDL1+ 0.05 0.02 0.01 *
Control Mild Severe
Positivity Fraction
L
r = 0.58
R² = 0.34
Fraction of CD103+ Neutrophils
Neutrophils density [Cells/mm²]

**Figure S3. Spatial mapping of human aGVHD reveals non-canonical immunological subtypes**

**(A)** Top: Total number of immune cells and their relative abundances (y-axis) across patients (x-axis) with a colonic biopsy (left) or duodenal biopsy (right). Bottom: The total number of non-immune cells and their relative abundances (y-axis) for each patient (x-axis). Below are shown clinical characteristics of the patients. **(B)** Bar plot displaying the distribution of immune subgroups, as determined per individual FOVs, across patients. The y-axis indicates the number of FOVs for each patient and the x-axis indicates the different patients with a duodenal biopsy (GVHD n=40). Bars are colored according to the classification of the FOVs into one of the immune subgroups shown in Fig. 4C. While control patients are rather homogeneous, GVHD shows increased intra-patient variability. **(C)** The number of patients (y-axis) that have either a single immune-subgroup or multiple subgroups (x-axis). **(D)** For each immune subgroup (x-axis), shown are the number of patients with at least one FOV in that subgroup. **(E)** Percentage of FOVs with clinically severe and mild GVHD (y-axis) in the Macs+CD8T subgroup compared to other immune subgroups (Chi-square, p value = 0.003). **(F)** Days after transplantation (y-axis) in the CD8+T enriched subgroup compared to other immune subgroups. Each dot represents a FOV from a patient. (Wilcoxon rank-sum, p value=0.001). **(G)** For each protein (rows), shown is the fraction of positive cells out of CD4+T cells in control, clinically mild, and clinically severe GVHD (columns). Patients with less than 20 CD4T cells were excluded from the analysis. Asterisks denote statistically significant changes relative to the control. P values are calculated by Wilcoxon rank-sum test followed by Benjamini-Hochberg p-value adjustment across rows, *P<0.05, **P<0.01, ***P<0.001). **(H-K)** Same as (G) for Tregs, plasma cells, macrophages and neutrophils, respectively. **(L)** Linear relationship between neutrophil density in the lamina propria and CD103+ neutrophils in GVHD patients. Linear regression is displayed in blue solid line with 95% confidence interval.

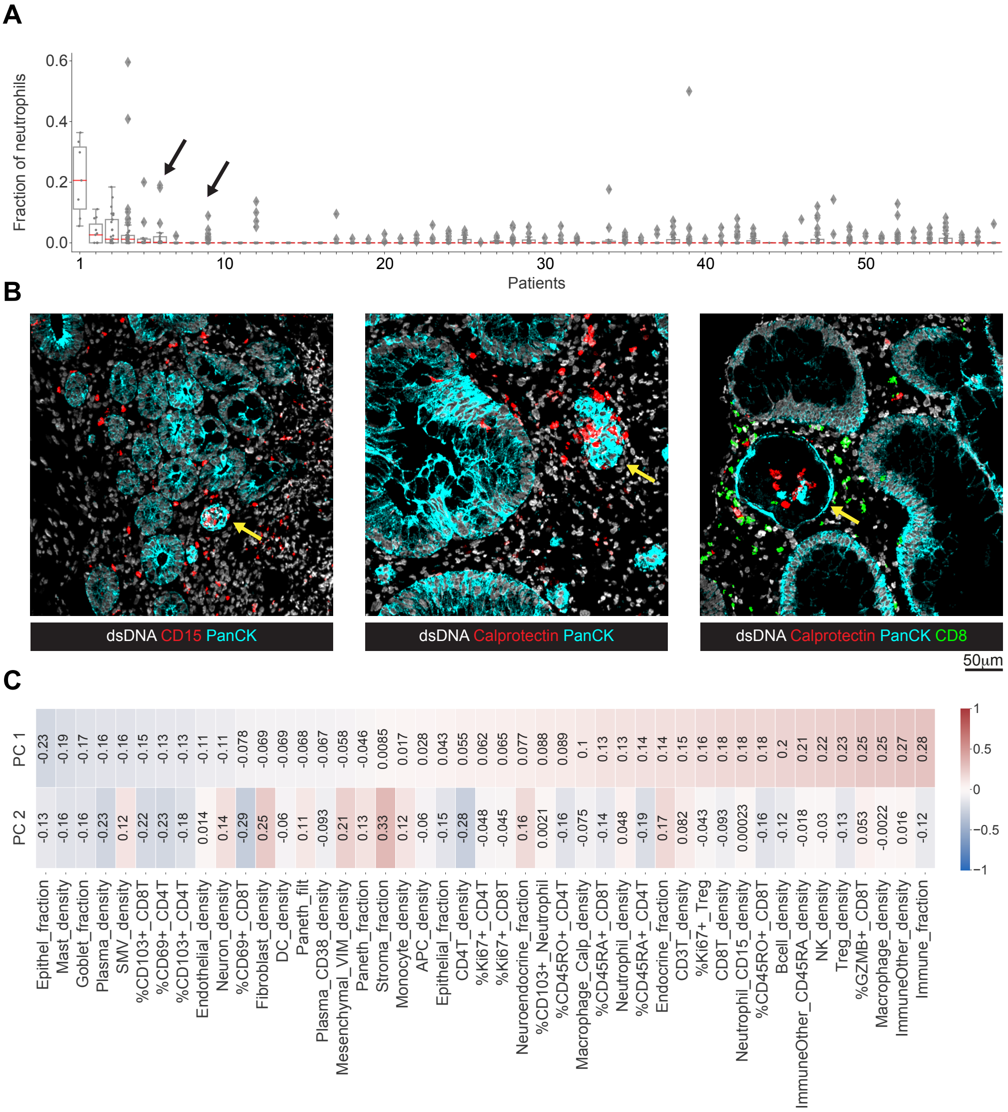


**Figure S4. Local immune microenvironments drive intra-patient heterogeneity**

**(A)** Individual crypt/villus objects were segmented using CellPose and dilated to quantify their local immune microenvironments. Shown is the fraction of Neutrophils out of all cells surrounding individual crypts. Box plots summarize all crypts in individual patients and depict the 25th and 75th percentiles. Dots represent individual crypts. The red line shows the median. **(B)** Images from three patients displaying a single epithelial object (indicated by an arrow) enriched with neutrophils (indicated by calprotectin or CD15). The objects also display a distorted morphology. In the rightmost image, the object is also surrounded by CD8T cells. **(C)** Loadings for the PCA plot in Fig. 7A. Features (rows) are ordered according to their contribution to PC1.
