## Supplementary material for "A spatial atlas of human gastro-intestinal acute GVHD reveals epithelial and immune dynamics underlying disease pathophysiology": Table S1

| Table 1. Patients' characteristics (59 biopsies)<br>Characteristics | | |
|---|---|---|
| **Age, median (range), yr** | 57 | ( 17-69) |
| **Gender, n (%)** | | |
| Male | 33 | (56) |
| Female | 26 | (44) |
| **Disease, n (%)** | | |
| AML | 32 | (54) |
| MDS | 27 | (46) |
| **Donor, n (%)** | | |
| MUD | 35 | (59) |
| Haplo | 8 | (14) |
| Sibling | 15 | (25) |
| MMUD | 1 | (2) |
| **Conditioning, n (%)** | | |
| Myeloablative | 15 | (25) |
| RIC | 44 | (75) |
| **Conditioning type, n (%)** | | |
| BuFlu | 34 | (58) |
| BuCy | 8 | (13) |
| FluMel | 7 | (12) |
| Other | 10 | (17) |
| **Added immunosuppression type, n (%)** | | |
| ATG | 36 | (61) |
| PTCy | 7 | (12) |
| None | 16 | (27) |
| **GVHD Prophylaxis** | | |
| CSA MMF | 46 | (78) |
| CSA MTX | 13 | (22) |
| **GVHD grade, n (%)** | | |
| 0 | 9 | (15) |
| 1 | 1 | (2) |
| 2 | 32 | (54) |
| 3 | 17 | (29) |
| 4 | 0 | (0) |
| **Stage GI, n (%)** | | |
| 0 | 10 | (17) |
| 1 | 24 | (41) |
| 2 | 18 | (30) |
| 3 | 7 | (12) |
| 4 | 0 | (0) |
| **Stage Liver, n (%)** | | |
| 0 | 53 | (90) |

| | | |
|---|---|---|
| 1 | 2 | (3) |
| 2 | 3 | (5) |
| 3 | 1 | (2) |
| 4 | 0 | (0) |
| **Stage Skin, n (%)** | | |
| 0 | 35 | (60) |
| 1 | 9 | (15) |
| 2 | 9 | (15) |
| 3 | 6 | (10) |
| 4 | 0 | (0) |
| **Pathological score GI, n (%)** | | |
| 0 | 25 | (42) |
| 1 | 18 | (31) |
| 2 | 10 | (17) |
| 3 | 6 | (10) |
| 4 | 0 | (0) |
| **Response to corticosteroids, n (%)** | | |
| Responder | 29 | (49) |
| Refractory | 23 | (39) |
| N/A | 7 | (12) |
| **Follow-up, n (%)** | | |
| Alive | 29 | (49) |
| Death | 30 | (51) |
| **Cause of death, n (%)** | | |
| NRM | 22 | (37) |
| Relapse | 8 | (14) |
| Alive | 29 | (49) |
| **Biopsy location, n (%)** | | |
| Colon | 19 | (32) |
| Duodenum | 40 | (68) |

Abbreviation

Acute myeloid leukemia (AML), Myelodysplastic syndrome (MDS), Matched unrelated donor (MUD), Mismatched unrelated donor (MMUD), Haploidentical (Haplo), Reduced intensity conditioning (RIC), Busulfan and Fludarabine (BuFlu), Busulfan and Cyclophosphamide (BuCy), Fludarabine and Melphalan (FluMel), Anti-thymocyte globulin (ATG), Post-transplantation Cyclophosphamide (PTCy), Cyclosporine A and Mycophenolate mofetil (CSA MMF), Cyclosporine A and Methotrexate (CSA MTX), Non relapse mortality (NRM), Not available (N/A)
