## Supplementary material for "A spatial atlas of human gastro-intestinal acute GVHD reveals epithelial and immune dynamics underlying disease pathophysiology": Table S2

Over night staining

| **Antibody** | **Clone** | **Vendor** | **Metal** | **dilution** |
|---|---|---|---|---|
| dsDNA | 35I9 | Ionpath | 89 | 25 |
| Collagen 1 | E8F4L | CST | 113 | 100 |
| CD206 | E2L9N | CST | 140 | 200 |
| CD103 alexa | EPR4166(2) | Abcam | 141 | 150 |
| Foxp3 | 236A/E7 | Abcam | 142 | 40 |
| CD4 | EPR6855 | Abcam | 143 | 40 |
| Granzyme B | D6E9W | CST | 144 | 75 |
| PD1 | d4w2j | CST | 145 | 50 |
| TCF1 | C63D9 | CST | 146 | 50 |
| CD45RA | HI100 | BD Pharmingen | 147 | 200 |
| CD45 | D9M8I | CST | 148 | 100 |
| PDL1 biotin | E1L3N | CST | 149 | 150 |
| CD69 | EPR21814 | Abcam | 150 | 200 |
| CD56 | MRQ-42 | Ionpath | 151 | 25 |
| CD31 | EP3095 | Ionpath | 152 | 100 |
| CD14 | D7A2T | CST | 154 | 50 |
| Mucin 2 | f2 | SANTA CRUZ | 155 | 100 |
| CD68 | D4B9C | Ionpath | 156 | 100 |
| IgA | EPR5367-76 | Ionpath | 157 | 200 |
| CD8 | C8/144B | Ionpath | 158 | 15 |
| CD3 | MRQ-39 | Ionpath | 159 | 15 |
| CD15 | BRA-4F1 | Abcam | 160 | 50 |
| CD45RO | UCHL1 | Ionpath | 161 | 25 |
| Calprotectin | MAC387 | Abcam | 162 | 200 |
| vimentin | D21H3 | Ionpath | 163 | 50 |
| Keratin | AE-1/AE-3 | BioLegend | 164 | 100 |
| CHGA | EPR22537-249 | abcam | 166 | 100 |
| CD20 | L26 | ionpath | 167 | 30 |
| Na/K | D4Y7E | CST | 168 | 50 |
| Ki-67 | 8D5 | CST | 169 | 50 |
| tryptase | EPR9522 | abcam | 170 | 400 |
| IDO-1 | EPR20374 | ionpath | 171 | 40 |
| HLA- DR,dq,dp | CR3/43 | abcam | 172 | 200 |
| CD209 | DCN46 | ionpath | 173 | 50 |
| CD38 | E7Z8C | ionpath | 174 | 50 |
| Lysozyme | SP329 | ionpath | 175 | 50 |
| HLA1 | EMR8-5 | ionpath | 176 | 50 |

1 hour staining

| Antibody | Clone | Vendor | Metal | dilution |
|---|---|---|---|---|
| SMA | SP171 | Abcam | 115 | 250 |
| alexa | polyclonal | Thermo Fisher Scientific | 141 | 125 |
| antibiotin | 1D4-C5 | BioLegend | 149 | 200 |
| CD163 | SMAD4 | SANTA CRUZ | 153 | 100 |
| E-cadhherin | 24E10 | CST | 165 | 50 |
